# A Zur-dependent regulatory RNA involved in maintaining zinc homeostasis in *Staphylococcus aureus*

**DOI:** 10.1101/2025.08.23.671911

**Authors:** Mathilde Charbonnier, Samuel Probst-Lotze, Hugo Racine, Jana N. Radin, Gustavo Rios-Delgado, Hannah M. Laster, Maximilian P. Kohl, Rafał Mazgaj, Marion Blum, Virginie Marchand, Johana Chicher, Stefano Marzi, Pascale Romby, Jai J. Tree, Kevin J. Waldron, Jeffrey M. Boyd, Julien Y. Dutheil, Thomas E. Kehl-Fie, David Lalaouna

## Abstract

Small regulatory RNAs (sRNAs) are key drivers of bacterial adaptation to environmental fluctuations, including iron and manganese restriction imposed by the host. This study explored the repertoire of sRNAs produced by the human pathogen *Staphylococus aureus* in response to metal limitation. Two sRNAs, S1077 and ZinS (RsaX20), regulated by zinc (Zn) availability, were identified. Further investigations revealed that, similar to the *cnt* operon from which it derives, S1077 synthesis is controlled by the transcription factors Zur and Fur. In contrast, *zinS* transcription is solely repressed by Zur. Amongst the ZinS targets are several Zn-dependent enzymes, such as the alcohol dehydrogenase Adh, whose synthesis is negatively regulated by ZinS. Loss of ZinS does not alter staphylococcal metal accumulation, suggesting a role in a Zn-sparing response. Remarkably, *zinS* also encodes a small peptide, ZinP. Genomic analysis suggests that the regulatory portion of ZinS emerged from the 3’ untranslated region of *zinP* in *S. aureus* and closely related species after horizontal gene transfer from phylogenetically distant organisms. All our findings demonstrate that sRNAs also facilitate bacterial adaptation to Zn limitation, and that genetic exchange and subsequent neofunctionalization have enabled *S. aureus* to adapt to metal-restricted environments.

## INTRODUCTION

In the environment and during infection, bacteria must cope with the scarcity of essential nutrients including transition metals (1). During infection, *Staphylococcus aureus* and other invaders experience acute iron (Fe), manganese (Mn), and zinc (Zn) starvation due to the active restriction of metal availability by the host, a defense known as nutritional immunity. Survival within the host and other metal-restricted environments requires adaptive changes that are mediated by a combination of proteinaceous and RNA-based regulatory mechanisms (2,3). While proteinaceous regulators are known to contribute to the ability of *S. aureus* and other microbes to adapt to changes in Zn availability, it is unknown if RNA-based regulatory mechanisms are used to respond to Zn limitation.

Zn is an essential trace element playing a pivotal role in numerous biological processes, such as enzymatic catalysis and protein structural stability (4). The host leverages this essentiality to control infection through the action of metal-sequestering proteins such as calprotectin (CP) (5,6). This S100 protein can be found at sites of infection at concentrations exceeding 1 mg/mL (7). CP can sequester a variety of transition metals, including Zn, which it binds with picomolar affinity (8–11). This tight binding allows CP to severely restrict Zn availability in culture and during infection (9,12). *S. aureus* and other microbes, including all ESKAPE pathogens, regulate their response to Zn limitation via Zur, a member of the ferric uptake regulator family (13). Zur is a zinc-dependent transcriptional repressor with a femtomolar sensitivity (14). Under zinc-replete conditions, Zn-bound Zur recognizes specific DNA sequences known as Zur boxes in the promoter region of target genes, thereby inhibiting their transcription. In the absence of Zn, Zur dissociates from DNA, leading to the expression of Zn transporters to enhance Zn acquisition and adaptations that enable the preservation of essential processes (15). In *S. aureus,* the Zur regulon encompasses genes involved in the synthesis of the Zn acquisition system (*acdA*, *acdBC-zur*, and the *cnt* operon), Zn-independent paralogs of the ribosomal proteins S14 (*rpsN2*) and L33 (*rpmG2*), and the Zn-binding GTPase ZagA (COG0523 subfamily of putative Zn chaperones) (16–18). Although the exact Zur regulon varies between species, a similar set of processes are controlled by this regulator.

While bacteria must cope with metal limitation, they must also contend with Zn intoxication within the host and the environment (19). In excess, Zn can inhibit the uptake of other metals and disrupt bacterial metabolism by displacing essential metal ions in enzymes (20,21), a phenomenon known as mismetallation. The dual nature of zinc necessitates mechanisms for the strict maintenance of its homeostasis in bacterial cells, leading bacteria, including *S. aureus*, to possess regulators and systems intended to prevent overaccumulation. Elevated levels of Zn are sensed by CzrA, a member of the ArsR family, which prevents the synthesis of the Zn efflux pump CzrB (22,23). Differing from Zur, apo-CzrA has a higher DNA binding affinity than Zn-bound CzrA, preventing repression of its regulon in the presence of zinc (24).

Numerous studies have shown the crucial role of regulatory small RNAs (sRNAs) in coordinating the bacterial response to Fe and Mn limitation (2,3). However, despite the critical importance of Zn, there are no sRNAs known to facilitate the bacterial response to Zn starvation. In *S. aureus*, IsrR and RsaC are involved in the Fe- and Mn-sparing response, respectively (25–28). The expression of both sRNAs is directly controlled by the respective metal-sensing transcription factors, Fur (Fe) and MntR (Mn) (29,30). This leads to the hypothesis that if sRNAs that modulate Zn utilization exist, they should be regulated by Zur. Leveraging this idea, this study found that metal starvation highly induces the expression of two additional putative regulatory sRNAs, S1077 and ZinS (formerly RsaX20 (31)). Subsequent investigation revealed that the synthesis of S1077 sRNA is induced by Zn and Fe depletion, similar to the *cnt* operon from which it derives. ZinS responds specifically to Zn starvation, being under the control of the transcription factor Zur. We also demonstrated that ZinS is a dual-function sRNA, coding for a small peptide, called ZinP. The regulatory part of ZinS most likely emerged from the 3’ untranslated region (3’UTR) of ZinP after horizontal gene transfer from phylogenetically distant organisms such as *Enterococcus* or *Streptococcus*. We then explored the Zur regulon and the ZinS targetome to unveil the participation of each protagonist. In particular, we observed that ZinS controls the synthesis of several Zn-dependent enzymes, including the alcohol dehydrogenase Adh, likely reducing Zn usage in response to Zn starvation.

## MATERIALS AND METHODS

### Strains, plasmids, and growth conditions

Bacterial strains and plasmids used in this study are listed in Table S1A. For plasmid construction, PCR fragments amplified by oligonucleotides indicated in Table S1B were digested with respective restriction enzymes and ligated into a similarly digested vector. To create the transcriptional reporter fusion, the *zinS* promoter and the first 19 nucleotides of *zinS* were PCR amplified using the indicated primers (Table S1B) and cloned into pAH5 (32). To generate the translational reporter, the 5’ UTR and the first 2 codons of ZinP were PCR amplified using the indicated primers (Table S1B) and cloned into pCN52 (33) under control of the pHELP promoter (34). Plasmids isolated from *E. coli* TOP10 (Thermo Fischer Scientific) were then transferred into indicated backgrounds. *E. coli strain* IM08B (35) was used as recipient strain for plasmid amplification and to efficiently transform *S. aureus* cells.

*S. aureus* strains were isolated on Tryptic Soy Broth (TSB) or blood agar plates. Overnight cultures were diluted 50-fold in NRPMI medium to induce metal starvation. To prepare NRPMI medium, 1% casamino acids (MP Biomedicals) and 5% Chelex-100 resin (Sigma-Aldrich) were added to Roswell Park Memorial Institute medium (RPMI; Sigma-Aldrich), then mixed for 6 hours at room temperature. The resulting medium was sterile-filtered and supplemented with metal ions at the following concentrations: 1 mM MgCl_2_, 100 µM CaCl_2_, 25 µM ZnCl_2_, 25 µM MnCl2, and 1 µM FeSO_4_ (Sigma-Aldrich). Cells were then grown with shaking at 37°C (180 rpm, 5:1 flask-to-medium ratio). Antibiotics were added as needed: 100 μg/mL of ampicillin, 10 μg/mL of chloramphenicol, and 10 μg/mL of erythromycin.

### Gene deletion

The deletion of the *zinS* (Δ*zinS*), *zur* (Δ*zur*), *fur* (Δ*fur*) and *mntR* (Δ*mntR*) genes in *S. aureus* HG001 strain was performed using the pMAD suicide vector (36). Upstream and downstream regions of the respective gene (≈600 nts) were amplified by PCR (Table S1B) and cloned into pMAD plasmid. The obtained plasmids were then transferred to the IM08B recipient strain and finally electroporated into *S. aureus* HG001. Mutant strains were selected as previously described (36). Cells were grown at restrictive temperature (44°C) and incubated at 28°C to favor a double crossover. The gene deletion was verified by PCR and sequencing. The same strategy was used to insert the MS2 sequence at the 5’end of *zinS* gene.

### Phylogenomic and molecular evolution analyses

Staphylococcal species from Barwinska-Sendra et al. (37) were selected for complete genome alignment. The reference genomes of each species were retrieved using the NCBI command line interface (CLI) tools. The genome of *Mamaliicoccus vitulinus* (formerly *S. vitulinus*) has 223 contigs and was discarded. All sequences annotated as plasmids were ignored. A pairwise distance matrix was generated using the alignment-free phylogeny reconstruction method Andi (38). A phylogenetic tree was subsequently generated with FastME (39) using default parameters. The distance-based phylogenetic tree was used as a guide tree for the cactus aligner (40) to produce a multiple genome alignment. A genome alignment in MAF format was then created after projection on the HG001 strain and further processed using MafFilter (41). The genome alignment was refined in blocks of maximum 10 kb using the MAFFT aligner (42). Synteny blocks containing all selected species and no duplicated sequences were retained. Site-filtering was applied by sliding a window of 10 nt by 1 nt. Any window containing more than two gaps in at least two species were discarded and the corresponding synteny block split. Block in synteny distant by less than 100 nt were then merged, and missing positions were replaced by N, resulting in masking of ambiguously aligned regions. Local phylogenetic trees were then inferred by removing blocks less than 500 nt long and splitting the remaining blocks into windows with an average size of 5 kb. A tree was built with the PhyML method (43), using the HKY85 (44) nucleotide substitution model and a four-classes gamma distribution of substitution rates, using the best tree from nearest neighbor interchange (NNI) and subtree pruning and regrafting (SPR) topology search algorithms. The 748 resulting local trees were given as input to the ASTRAL-IV (45) method from the ASTER package (https://github.com/chaoszhang/ASTER) in order to get a phylogeny, which was subsequently rooted using the MAD method (46). The resulting phylogeny slightly differed from the alignment-free guide tree and was consistent with that in Barwinska-Sendra et al. (37). It was used as a new guide tree to the cactus aligner to generate a new genome alignment. This final alignment was used to extract the *zinS* locus sequence.

A blast nucleotide database was created with all complete genomes to look at possible paralogs of the *zinS* locus. The ZinP amino acid sequence was used as a query for the tblastn program. All matching sequences with an E-value lower than 1e-6 were extracted and aligned with the ClustalO aligner (47). The alignment was converted to PAML input file format using the Bio++ Program Suite (48); the HG001 sequence was discarded for this analysis, as it is identical to the reference sequence for *S. aureus*. The codeml program from the PAML 4.10.7 package was used to fit models of codon sequence evolution (49). Several models allowing for different selective pressure along the tree were serially tested to select the best model fit.

### Small RNA-sequencing

Total RNA was extracted from cells grown in NRPMI supplemented with 1 mM MgCl_2_ and 100 µM CaCl_2_ or RPMI + 1% CA (Control) and harvested at OD_600nm_ = 1.5. RNA was treated with DNase I (Roche) according to the manufacturer’s recommendations. RNA was selected using the PippinPrep instrument (80-300 nts; Sage Science). Note that the unexpected detection of large regulatory RNAs like RsaC is due to the presence of shorter fragments. RNA quantity and quality were assessed using the 2100 Bioanalyzer (Agilent). The cDNA library was obtained using the NEBNext® Small RNA Library Prep Set for Illumina®. and sequenced with the NextSeq 2000 system (Illumina). The ligation of adaptors to unfragmented RNA was performed after treatment with RppH (NEB). Differential gene expression analysis was performed using the READemption pipeline (50). Reads were mapped to sRNA features only. Small RNA boundaries were defined using the *S. aureus* strain RN1 annotation from the *Staphylococcus* regulatory RNA database (51). The RN1 sRNA annotation was transferred to the HG001 genome (NZ_CP018205.1) (52) using RATT (53) and the Strain.global setting. Data were visualized using VolcaNoseR (54). Results are representative of three independent experiments.

### Gel retardation assays

Transcription start sites were determined according to Koch et al. (55) with the exception of *zinS*. An arbitrary value of -200 nt was used when no clear +1 was observed. PCR fragments containing T7-5’*adh* (from nt -276 to +564), T7-3’*adh* (from nt +540 to +1115), T7-*isaB* (full-length; from nt -39 to +589), T7-*tdh* (from nt -67 to +662), T7-*hslO* (from nt -200 to +676), T7-*mntR* (full-length; from nt -20 to +713), T7-*sepF* (from nt -200 to +568), and BamHI-digested pUT7 plasmids containing 5’*aur* (from nt -40 to +745) or 3’*aur* (from nt +718 to 1660) were used as DNA templates for *in vitro* transcription with homemade T7 RNA polymerase. T7-*zinS* was cloned into pJET1.2/blunt plasmid (ThermoFisher). RNAs were finally purified and radiolabeled when required (25). 5’-radiolabeled ZinS (20,000 cpm/sample) and the aforementioned cold RNAs were separately denatured at 90°C in GR-buffer (20 mM Tris-HCl pH 7.5, 60 mM KCl, 40 mM NH4Cl, 3 mM DTT), immediately cooled on ice for 1 minute, and incubated at room temperature for 15 minutes in the presence of 10 mM MgCl_2_. The renatured RNAs were then mixed and incubated at 37°C for 15 minutes. The specificity of the interaction was tested by competition with cold ZinS and yeast tRNA when indicated. Finally, the samples were loaded onto a 6% polyacrylamide gel under non-denaturing conditions. Gels were exposed to Fuji X-ray films and developed in an Optimax X-ray film processor. The results are representative of two independent experiments.

### Northern blot analysis

Samples were collected at specified OD_600nm_. After centrifugation, the bacterial pellets were resuspended in RNA Pro Solution from the FastRNA Pro Blue kit (MP Biomedicals). Cell lysis was achieved using the FastPrep homogenizer (MP Biomedicals). Total RNA extraction was then carried out according to the manufacturer’s recommendations. 5 µg of total RNA were loaded onto a 6% polyacrylamide gel and electrophoresed at 120 V. RNA was then transferred to a Hybond N+ nitrocellulose membrane (GE Healthcare Life Sciences). Membranes were hybridized with specific digoxygenin (DIG)-labeled probes, which were prepared using the DIG RNA Labeling Kit (Roche). Luminescent detection was performed using Anti-Digoxigenin-AP, Fab fragments and CDP-Star (Roche). The RiboRuler Low Range RNA Ladder (Thermo scientific) was used to determine the length of S1077 related bands. Results represent at least two independent experiments.

### Calprotectin growth assays with transcriptional and translational fusions

CP growth assays were performed as described previously (56). Briefly, overnight cultures (grown 16-18 h in 5 ml of NRPMI supplemented with 1 mM MgCl_2_, 100 µM CaCl_2_ and 1 µM FeSO_4_ in 15 ml conical tubes at 37°C on a roller drum) were used directly and diluted 1:100 into 96-well round-bottom plates containing 100 µl of growth medium (38% TSB and 62% calprotectin buffer (20 mM Tris pH 7.5, 100 mM NaCl, 3 mM CaCl_2_)) supplemented with 1 µM MnCl_2_ and 1 µM ZnSO_4_ in presence of varying concentrations of CP. For all assays, the bacteria were incubated with orbital shaking (180 rpm) at 37°C and growth was measured by assessing optical density (OD_600nm_) every 2 hours. Prior to measuring optical density, the 96-well plates were vortexed. Expression of *zinS* was assessed by measuring yellow fluorescence (excitation and emission wavelengths of 505 nm and 535 nm, respectively), normalized to OD_600nm_, as previously described (12). ZinP translation was assessed by measuring green fluorescence (excitation and emission wavelengths of 485 and 528, respectively), normalized to OD_600nm_. Data correspond to the mean of three independent experiments ± SD.

### Alcohol dehydrogenase (ADH) activity assay

Overnight cultures were diluted to an OD_600nm_ of 0.05 in 3 mL of fresh TSB medium and cells were cultured with shaking (250 rpm) at 37°C for 16 hours. 1 mL of cell suspension was pelleted by centrifugation and washed with 0.05 M phosphate buffer pH 8. Cell pellets were resuspended in 100 µL of phosphate buffer containing 20 µg each of DNase I (ITW Reagents) and lysostaphin (AMBI). After 1 hour of incubation at 37°C, cell debris was separated from lysates by centrifugation. The reaction mixture (1 mL) contained potassium phosphate buffer pH 7.5 (83.35 mM), 0.666 mM NAD+, 150 mM ethanol, and 40 µL of cell-free lysate. The formation of NADH (ε340 = 6220 M^-1^ cm^-1^) was followed spectrophotometrically at A_340nm_ (57) for three minutes using a Cary 60 UV-Vis Spectrophotometer (Agilent). Protein concentrations of the cell-free lysates were determined by Bradford protein colorimetric assay (Bio-Rad Protein Assay Dye Reagent Concentrate) modified for a 96-well plate. Data correspond to the mean of three independent experiments ± SD.

### Primer extension assays

Primer extension assays were conducted using 30 µg of DNase I (Roche)-treated total RNA, incubated with a 5’-radiolabeled oligonucleotide (Table S1B), 2.5 mM dNTPs, and 4 units of AMV reverse transcriptase (NEB). The RNA template was removed by adding 3 µL of 3 M KOH and 20 µL of Destroy Buffer (50 mM Tris–HCl pH 8.0, 0.5% SDS, 7.5 mM EDTA) and incubating for 3 minutes at 90°C, followed by 1 hour at 37°C. The samples were then precipitated and run on a denaturing 10% polyacrylamide gel alongside a sequencing ladder. The sequencing ladder was produced using a DNA template with ddNTPs added to terminate the reaction catalyzed by 2 units of Vent (exo-) DNA polymerase (NEB). PCR was performed with the following conditions: 1 minute at 95°C, 1 minute at 52°C, and 1 minute at 72°C for 25 cycles. The results are representative of two independent experiments. Note that the presence of additional bands can be attributed to premature termination of reverse transcription (e.g., strong secondary structure or presence of RNA modifications).

### Toeprinting assays

Toeprinting assays were performed as described by Fechter et al. (58). 1 pmol of *in vitro* transcribed ZinS sRNA and 4 pmol of radiolabelled primer were mixed in a total volume of 9 µL, denatured at 90 °C, re-folded on ice for 1 min and incubated at room temperature for 5 min before the addition of 1 µL of TP10X+ buffer (100 mM MgCl_2_, 200 mM Tris-HCl pH 7.5, 600 mM NH_4_Cl and 10 mM DTT). In parallel, purified *S. aureus* 30S ribosomes were adjusted to 2 µM in TP1X+ buffer and pre-incubated at 37°C for 15 min. Next, 2 µL of mRNA/primer, 2 µL of 30S, 1 µL of 50 mM MgCl_2_ and 1 µL of TP10X-buffer (200 mM Tris-HCl pH 7.5, 600 mM NH_4_Cl and 10 mM DTT) were mixed in a total volume of 10 µL. Increasing concentrations of ZinS (50 - 400 nM) were added when indicated. Samples were incubated for 10 min at 37°C before the addition of 1 µL of 20 μM initiator tRNA^fMet^. After 5 min at 37°C, 2 µL of RT mix (0.2 mM dNTPs and 1 U/µL AMV RT in AMV RT buffer 1x; Promega) were added. After 30 min at 37°C, cDNA was extracted using PCI (Roth) and precipitated by addition of 2.5 volumes absolute ethanol and 0.1 volume of 3 M NaOAc pH 5.2. Pellets were washed, air dried and resuspended in urea blue loading buffer. Radioactivity (cpm) was measured and equal amounts resolved on a 10 % denaturing polyacrylamide gel. The results are representative of two independent experiments.

### 3’RACE

Total RNA was treated with DNase I (Roche). A ligation step was performed to add a 3’-blocked adaptor to the 3’ end of RNA (59). 4 µg of total RNA was incubated with 10 µM adaptor at 90°C for 2 minutes, followed by cooling on ice. Then 1X T4 RNA ligase buffer (NEB), 1 mM ATP, 100 U RNasin Plus (Promega) and 20 U T4 RNA ligase 1 ssRNA (NEB) were added and the ligation reaction was performed overnight at 16°C. RT-PCR was performed using ZinS- and adaptor-specific oligonucleotides and the One-Step RT-PCR Kit (Qiagen). Nested PCR was used to increase specificity and yield. The PCR products were then cloned into the pJET 1.2 vector. Finally, the resulting plasmids were sequenced (Eurofins GATC).

### ICP-OES analysis

Samples were collected at OD_600nm_ = 1. After centrifugation, the pellets were washed twice with 0.1 mM EDTA DPBS and twice with DPBS. Dry cell pellets were digested with concentrated nitric acid (65%, Merck) and heated at 70°C for 16 h. Samples were then diluted 10-fold into a 1% nitric acid solution containing 50 µg/L iridium (internal standard). Calibration curves were generated using matrix-matched standard solutions containing Mn, Fe, Zn, Cu, Mg, Ca and S. All standards and samples were analyzed by inductively coupled plasma optical emission spectrometry (ICP-OES) on a Thermo iCAP PRO instrument (RF power 1250 W, with nebulizer gas flow 0.5 L/min, auxiliary gas flow 0.5 L/min, and cool gas flow 13.5 L/min argon). The concentration of each element was determined by comparing to the standard curve using Qtegra ISDS Software (Thermo), and normalized to the intracellular level of sulfur (ppb). Data correspond to the mean of four independent experiments ± SD.

### Statistical analysis

Statistical analyses were performed using a two-way ANOVA followed by a Sidak’s multiple comparisons test or a Mann–Whitney test (Prism software). All statistical details of the experiments can be found in the Figure legends.

## RESULTS

### Identification of sRNAs induced by metal limitation

Knowing that regulatory RNAs are involved in the iron- and manganese-sparing responses in *S. aureus* (25–28), we hypothesized that sRNAs must also exist for other metals, including zinc. Therefore, *S. aureus* was grown in Chelex-treated RPMI + 1% casamino acids supplemented with 1 mM MgCl_2_ and 100 µM CaCl_2_ (NRPMI; metal-depleted condition) and RPMI + 1% casamino acids (control condition). Size-selected RNAs (80-300 nts) were extracted and adaptors were ligated to the unfragmented, RppH-treated RNAs. The RNAs were then subjected to differential RNA sequencing analysis (Figure 1). As expected, the synthesis of IsrR and RsaC was triggered in response to metal starvation. The modest increase of RsaC (1,116 nts) level compared to IsrR (174 nts) is likely explained by the size selection carried out prior to sequencing. In total, 42 RNA transcripts were up-regulated (e.g., S1077, ZinS (RsaX20), RsaOW2) and 42 were down-regulated (e.g., RsaI, RsaH, SprA2, SprC) in response to metal depletion (log_2_FC>|1|; -log10(padj)>1.3). The whole list is presented in Table S2. S1077 and ZinS (RsaX20) are remarkably responsive to metal starvation (log_2_FC = 7.2) and both are located near genes that facilitate staphylococcal resistance to host-imposed Zn and Mn starvation. S1077 is present in the 5’ UTR of the *cnt* operon, responsible for the biosynthesis and transport of the metallophore staphylopine (12,16). The presence of *zinS* (*rsaX20*), immediately adjacent to the gene encoding Mn-independent phosphoglycerate mutase GpmA (60), suggests it could be involved in coordinating the staphylococcal response to Mn availability. However, we also noticed the presence of a putative Zur-box upstream of ZinS (RsaX20), suggesting a role in Zn homeostasis.

**Figure 1.**
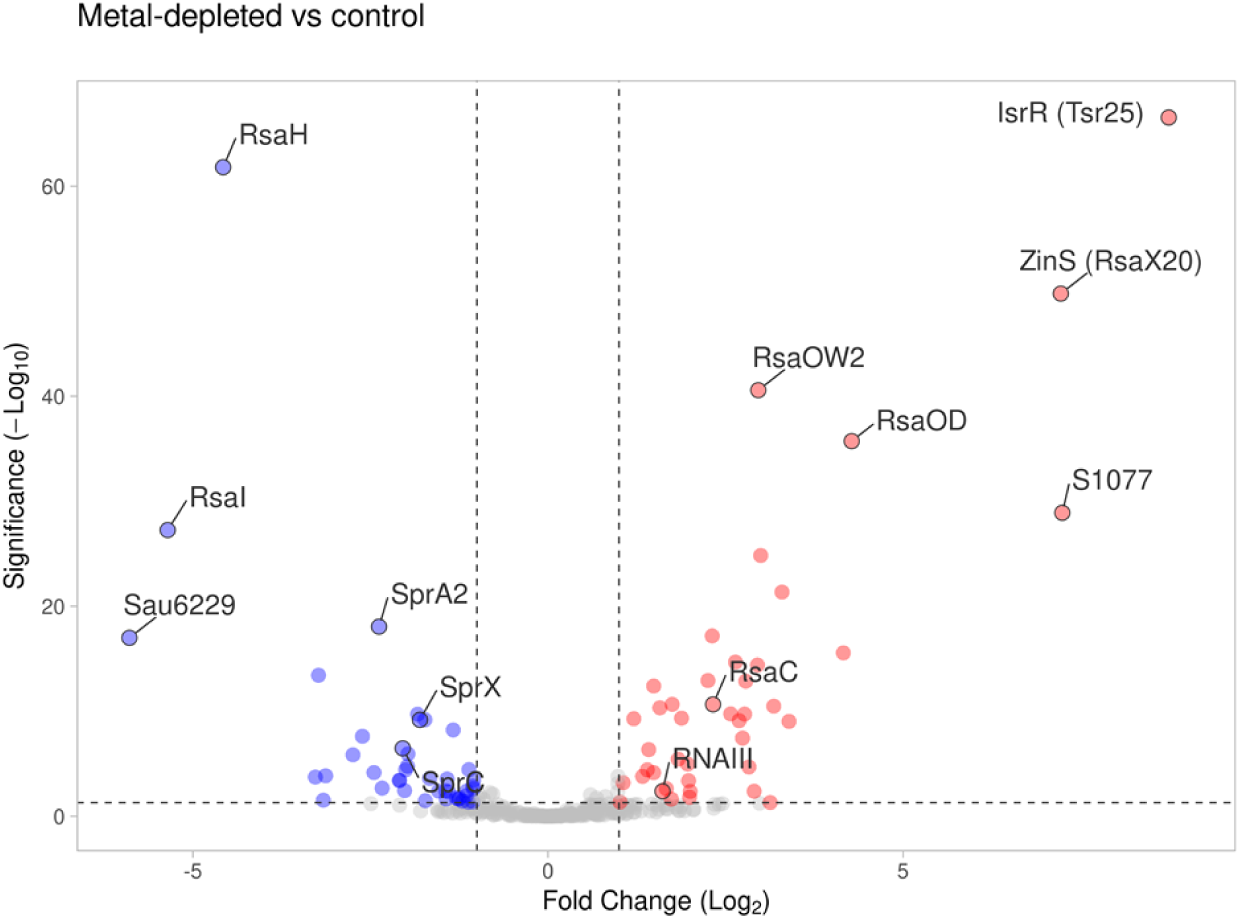
ZinS (RsaX20) and S1077 sRNAs are highly induced by metal limitation. sRNA sequencing analysis of size-selected RNAs (≈80-300 nts) extracted from *S. aureus* WT cells grown in NRPMI medium supplemented with 1 mM MgCl2 and 100 µM CaCl2 (metal-depleted condition) and RPMI + 1% casaminoacids (control condition). Data were visualized using VolcaNoseR (Goedhart et al., 2020) and are representative of three independent experiments. The following thresholds were applied: -log10(padj)>1.3; log2FC>|1|.

### S1077 is a 5’UTR-derived sRNA induced in response to Zn and Fe limitation

Given its integration in the 5’UTR of the *cnt* operon, we first determined S1077 boundaries using primer extension assays (Figures 2A and S1A) and rapid amplification of cDNA ends (3’RACE) (Figure S1B). The obtained results fit with the 5’ and 3’ ends identified by our sRNA-seq analysis (Figure S1C), suggesting that S1077 is a 220-nt-long sRNA (Figure 2B). We then showed that the 5’ end of S1077 is tri-phosphorylated, thereby protecting S1077 from degradation by the Terminator 5’-Phosphate-Dependent Exonuclease (Figure S1D). This indicates that S1077 and the *cnt* operon share the same promoter. Its secondary structure was finally predicted based on collected data (Figure 2C).

**Figure 2.**
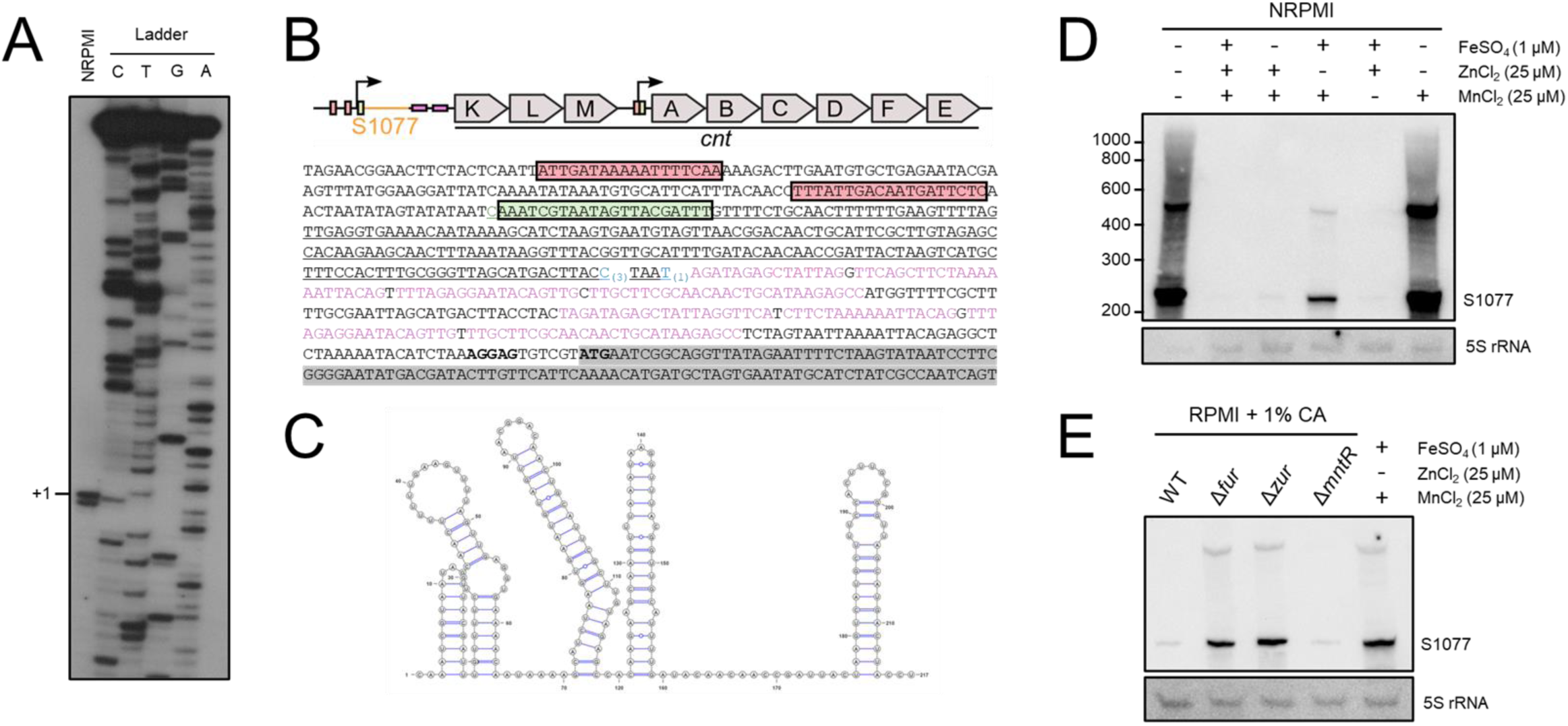
S1077 is a 220nt-long sRNA encoded in the 5’UTR of the *cnt* operon. A. Primer extension assay performed on total RNA extracted in NRPMI medium supplemented with 1 mM MgCl2 and 100 µM CaCl2 (OD600nm = 1) using a 5’-radiolabelled oligonucleotide complementary to the S1077 sequence. The primer is located at nucleotides +50 to +71 (from the +1 of S1077). The determined transcription start of S1077 is indicated by +1. C, T, G, A: sequencing ladders. Data are representative of two independent experiments. See also Figure S1A. B. Genomic organization of the *cnt* operon. The DNA sequence of the promoter and the 5’ untranslated region of the *cnt* operon is shown below. The red and green squares represent the Fur and Zur boxes, respectively. The sequence of S1077 is underlined. The start site (+1) is shown in green. Putative 3’ ends are shown in blue, with numbers in brackets representing the number of clones obtained. The two repeated sequences identified by Fojcik et al. (2018) are indicated in purple. The coding sequence of *cntK* is highlighted in grey. The Shine-Dalgarno sequence and the start codon (ATG) are in bold. C. Secondary structure of S1077 sRNA predicted using RNAalifold (Bernhart et al., 2008) and visualized using vaRNA (Darty et al., 2009). D. Northern blot analysis of S1077 *s*RNA levels. Total RNA was extracted from HG001 WT cells grown in NRPMI medium supplemented with 1 mM MgCl2, 100 µM CaCl2, ± 25 µM ZnCl2, ± 25 µM MnCl2 and ± 1 µM FeSO4. Cells were harvested at OD600nm = 1. 5S rRNA was used as loading control. RNA migrated along with a RiboRuler Low Range RNA Ladder (Thermo scientific). Results are representative of two independent experiments. E. Northern blot analysis of S1077 *s*RNA levels in HG001 WT, Δ*fur*, Δ*zur* and Δ*mntR* backgrounds in RPMI + 1% casamino acids medium. Cells were harvested at OD600nm = 1. The -Zn, -Fe and -Mn sample from Figure 2D was used as control. See also Figure S1.

To check whether S1077 sRNA is regulated similarly to the *cnt* operon, levels of S1077 were assessed using Northern blots. Total RNA was extracted from HG001 WT strain grown in NRPMI medium supplemented with Fe, Zn or Mn (Figures 2D and S1E). Using a S1077-specific probe, we observed two clear bands. The lower, main band corresponds to the expected length (220 nts), while the upper band (≈500 nts) potentially contains two repeated sequences, previously identified between S1077 and *cntK* (16) (Figure 2B). Although an increase of their levels was observed in the absence of Zn, the maximum was reached when depleting Zn and Fe. In agreement with this observation, we noticed an increase in S1077 levels in Δ*zur* and Δ*fur* mutant strains grown in RPMI + 1% casamino acids (Figure 2E).

All in all, our data support the involvement of the Zur and Fur transcription factors in the control of S1077 synthesis, similar to the *cnt* operon (12,16).

### ZinS (RsaX20) and the encoded peptide ZinP are induced in response to Zn limitation

ZinS (RsaX20) was identified in a high-throughput screen of intergenic regions with predicted RNA structure (31), but little is known about its boundaries, regulation or function. We initially determined the boundaries of this putative regulatory RNA using primer extension assays and 3’RACE. The identified (Figure 3A) and characterized transcription start site (5’PPP; Figure S2A) of ZinS (Figures 3A and S2A) was consistent with both the present sRNA-seq analysis, and the previous dRNA-seq analysis from Mediati et al. (61) (Figures S2B). Despite the substantial variability observed at the 3’ end (Figures 2B and S2C), the longest observed form of ZinS (RsaX20), which ends with a well-defined, Rho-independent terminator containing a polyU stretch (Figures 3B-C), was confirmed by a previous Term-seq analysis (61) (Figures S2B). Remarkably, it corresponds to a bidirectional transcription terminator (predicted using the TransTermHP algorithm (62)), suggesting that the 3’ ends of *zinS* (*rsaX20*) and *gpmA* may overlap. Using 3’RACE assays, we confirmed that both share 50 nts (Figure S2C-D). In total, this indicated that ZinS (RsaX20) is a 274 nt long sRNA.

**Figure 3.**
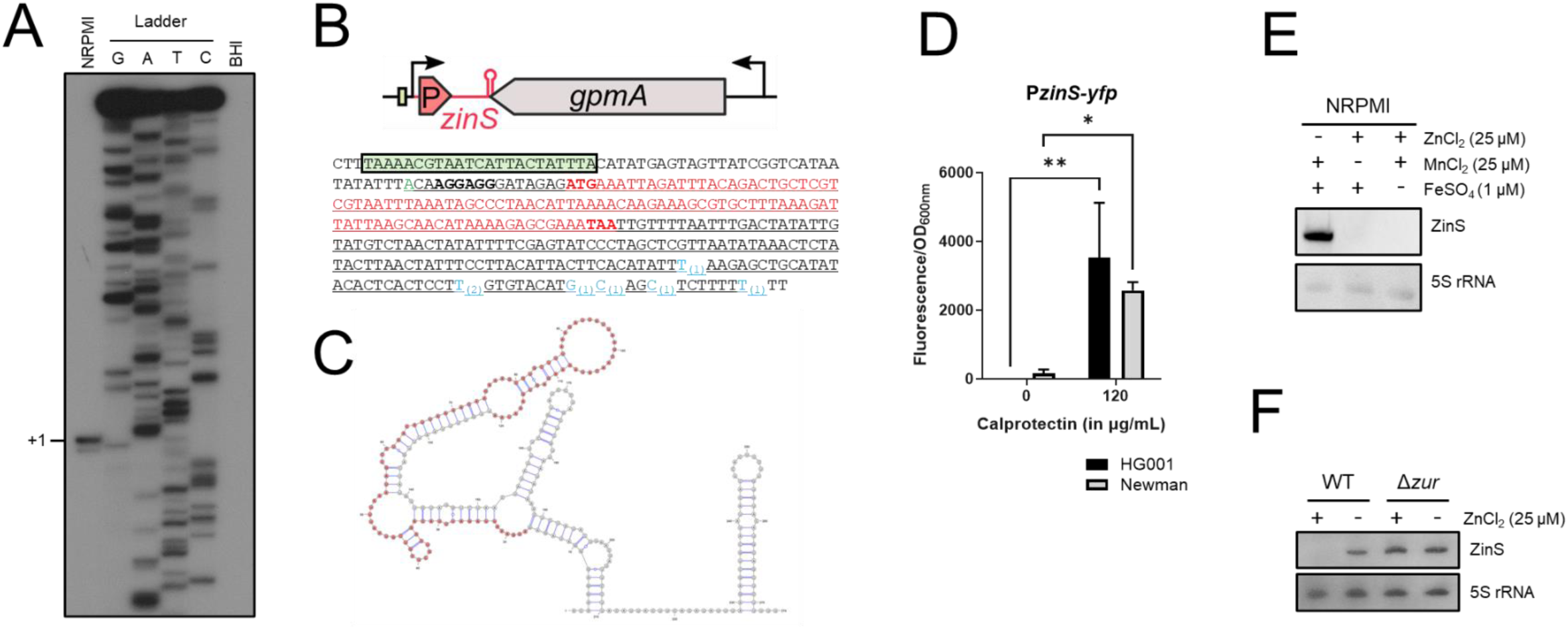
ZinS is a 274nt-long sRNA repressed by the Zn-dependent transcription factor Zur in *S. aureus*. A. Primer extension assay performed on total RNA extracted in NRPMI medium supplemented with 1 mM MgCl2 and 100 µM CaCl2 (OD600nm = 1) using a 5’-radiolabelled oligonucleotide complementary to the *zinS* sequence. The primer is located at nucleotides +44 to +64 (from the +1 of ZinS). The determined transcription start site of ZinS is indicated by +1. G, A, T, C: sequencing ladders. RNA extraction from cells grown in BHI was used as a negative control. Data are representative of at least two independent experiments. B. Genomic organization of the *zinS* gene. The DNA sequence is shown below. The start site (+1) is shown in green. Putative 3’ ends are shown in blue, with numbers in brackets representing the number of clones obtained. The coding sequence is highlighted in red (see Figure 4). The Shine-Dalgarno sequence (AGGAGG), the start codon (ATG) and the stop codon (TAA) are in bold. The Zur box is squared (green). C. Secondary structure of ZinS sRNA predicted using RNAalifold (Bernhart et al., 2008) and visualized using vaRNA (Darty et al., 2009). The coding sequence is shown in red. D. HG001 (black) and Newman (gray) cells containing the P*zinS-yfp* transcriptional fusion were grown in the presence or absence of calprotectin (120 µg/mL). The ratio of fluorescence/OD600nm was determined after 8 h of growth. Data shown are the combined results from three independent experiments ± SD. A two-way ANOVA with Šídák’s multiple comparisons test was performed using Prism software (*, P-value<0.05; **, P-value<0.005). E. Northern blot analysis of ZinS RNA levels in NRPMI medium supplemented with 1 mM MgCl2, 100 µM CaCl2, ± 25 µM ZnCl2, ± 25 µM MnCl2 and ± 1 µM FeSO4. HG001 WT cells were harvested at OD600nm = 1. 5S rRNA was used as loading control. Results are representative of two independent experiments. F. Northern blot analysis of ZinS RNA levels in HG001 WT and Δ*zur* backgrounds in NRPMI medium supplemented with 1 mM MgCl2, 100 µM CaCl2, 25 µM MnCl2, 1 µM FeSO4 ± 25 µM ZnCl2. See (E) for more details. See also Figures S2 and S3.

Given the uncertainty regarding the inducing metal, the transcription of ZinS (RsaX20) was first evaluated in the presence of calprotectin, observing induction at 120 µg/mL in both HG001 and Newman backgrounds (Figure 3D). To determine which metal ZinS (RsaX20) responds to, we monitored the level of ZinS (RsaX20) via Northern blot following HG001 WT strain growth in NRPMI medium supplemented with Zn, Mn, or Fe. A ZinS (RsaX20)-specific signal was only observed in media lacking Zn (Figure 3E). The same result was obtained using a promoter fusion (Figure S3A). We then deleted the genes encoding the two Zn import systems to block Zn acquisition, and drastically reduce intracellular Zn concentrations. Loss of the Adc and staphylopine-Cnt Zn import systems, but not the MntABC and MntH Mn importers, led to a significant increased transcription of the sRNA following growth in the presence of CP in Newman strain (Figures S3B-C). Therefore, *zinS* expression is strongly dependent on the intracellular concentration of Zn. The presence of a predicted Zur box suggests that ZinS responds directly to Zn availability via Zur. Consistent with this model, the loss of Zur led to constitutive expression of ZinS in metal replete medium (Figures 3F and S3D). Considering the Zn-specific regulation, we renamed this sRNA ZinS, for <u>Zin</u>c-responsive <u>s</u>mall RNA.

A short open reading frame encoding a small peptide of 34-amino acids (initially referred to as sRNA34; HG001_02453 or SAOUHSC_02702 in NCTC8325 background) was detected within ZinS using ribosome profiling (63). This peptide was renamed ZinP (Figures 3B and 4A). Toeprinting assays confirmed binding of the 30S ribosomal subunit to the Shine-Dalgarno (SD) sequence upstream of the start codon (Figure 4B), consistent with our previous findings using Retapamulin-enhanced ribosome profiling (Figure S4) (63). To assess the translational regulation of ZinP *in vivo*, we constructed a ZinP+6-GFP translational fusion. Upon exposure to calprotectin, we observed a strong increase in fluorescence in both HG001 and Newman backgrounds (Figure 4C), indicating that ZinP is produced under zinc-limiting conditions. Intriguingly, *zinP* homologous genes are frequently located in the vicinity of genes containing ZnuA/ZinT domains, potentially linking its function to Zn homeostasis (Figure S5). Although the role of ZinP remains to be elucidated, its predicted structure (Figure 4A) shows notable similarity to the MntS peptide in *E. coli*, which inhibits Mn efflux by obstructing the channel of the MntP exporter (64). To explore a potential analogous function for ZinP, we overexpressed it using a strong and constitutive promoter (P*blaZ*) and observed a substantial growth delay in the presence of toxic zinc levels (Figure S6A).

**Figure 4.**
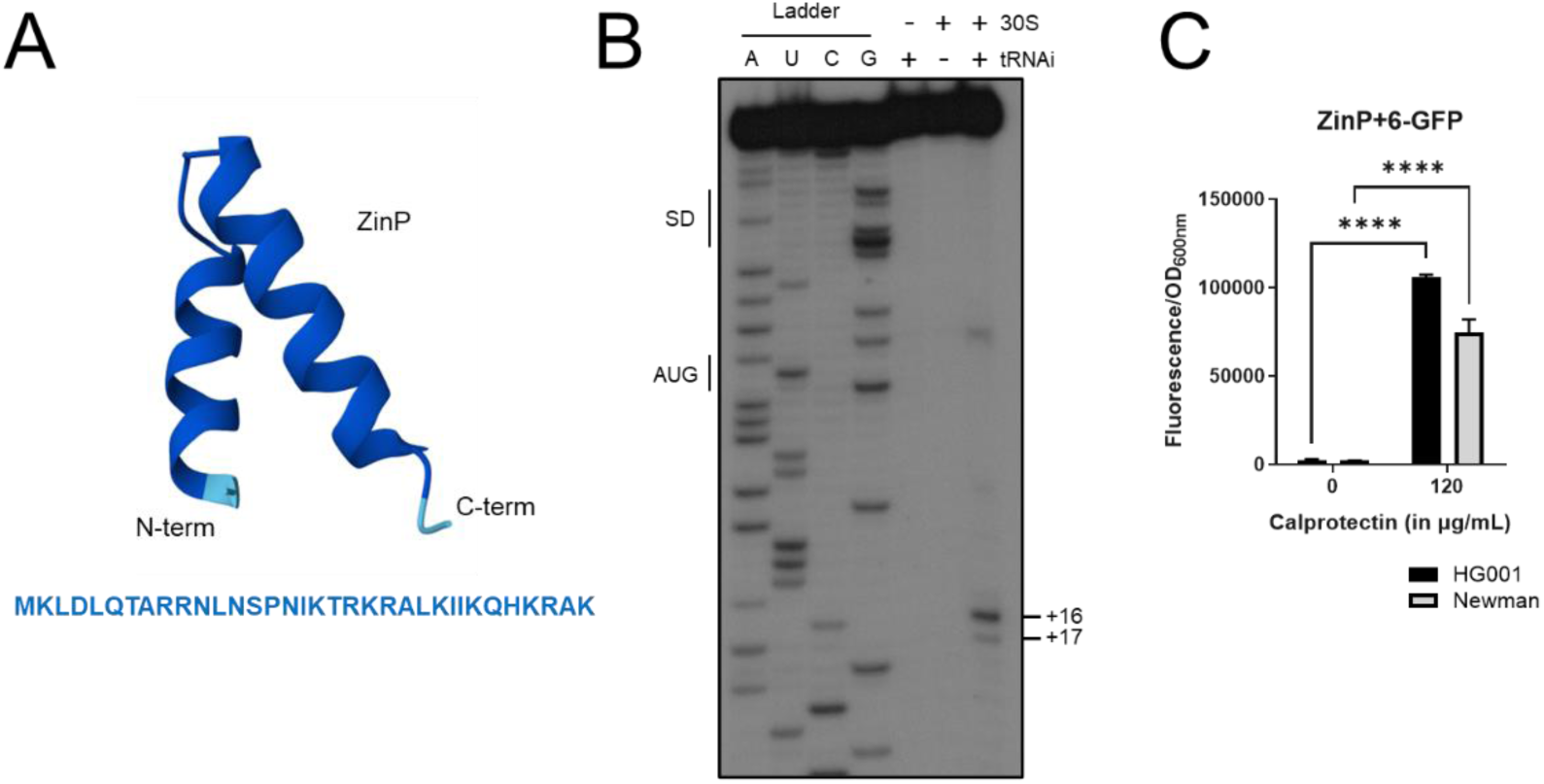
ZinS sRNA codes for the 34-amino-acid peptide ZinP. A. Amino acids sequence and ZinP structure predicted using AlphaFold3 (default parameters; Abramson et al., 2024). B. Toeprinting assay monitoring the formation of a ternary ribosomal initiation complex comprising *S. aureus* 30S subunit, initiator tRNA^fMet^ (tRNAi) and ZinS sRNA (50 nM). A, U, C, G: sequencing ladders; SD: Shine-Dalgarno sequence; AUG: start codon; +16/+17: toeprinting signal. Data are representative of two independent experiments. C. HG001 (black) and Newman (gray) cells containing the ZinP+6-GFP translational fusion were grown in the presence or absence of calprotectin (120 µg/mL). The ratio of fluorescence/OD600nm was determined after 8 h of growth. Results are representative of three independent experiments ± SD. A two-way ANOVA with Šídák’s multiple comparisons test was performed using Prism software (****, P-value<0.0005). See also Figure S4.

Altogether, these results indicate that ZinS and the small peptide ZinP that it encodes are synthesized specifically in response to zinc starvation and are under the direct control of Zur.

### ZinS and ZinP peptide emerged via horizontal gene transfer and neofunctionalization

An initial analysis suggested that the prevalence of ZinS is restricted to a subgroup of the staphylococci. Although ZinS is only highly conserved in *S. aureus* and closely related strains such as *S. schweitzeri* (96% sequence identity) and *S. argenteus* (97%), the 5’ part corresponding to the ZinP protein-coding region is conserved in other staphylococci species (Figure 5A). Therefore, using a multiple genome alignment of staphylococcal species, the underlying species phylogeny was reconstructed and orthologous regions of the *zinP* locus identified. This analysis places its origin in the common ancestor of *S. aureus* and *S. simulans* (Figure 5B). A tblastn search also revealed the occurrence of a ZinP protein at a distinct location in *Mammaliicoccus lentus*, albeit without a start codon.

**Figure 5.**
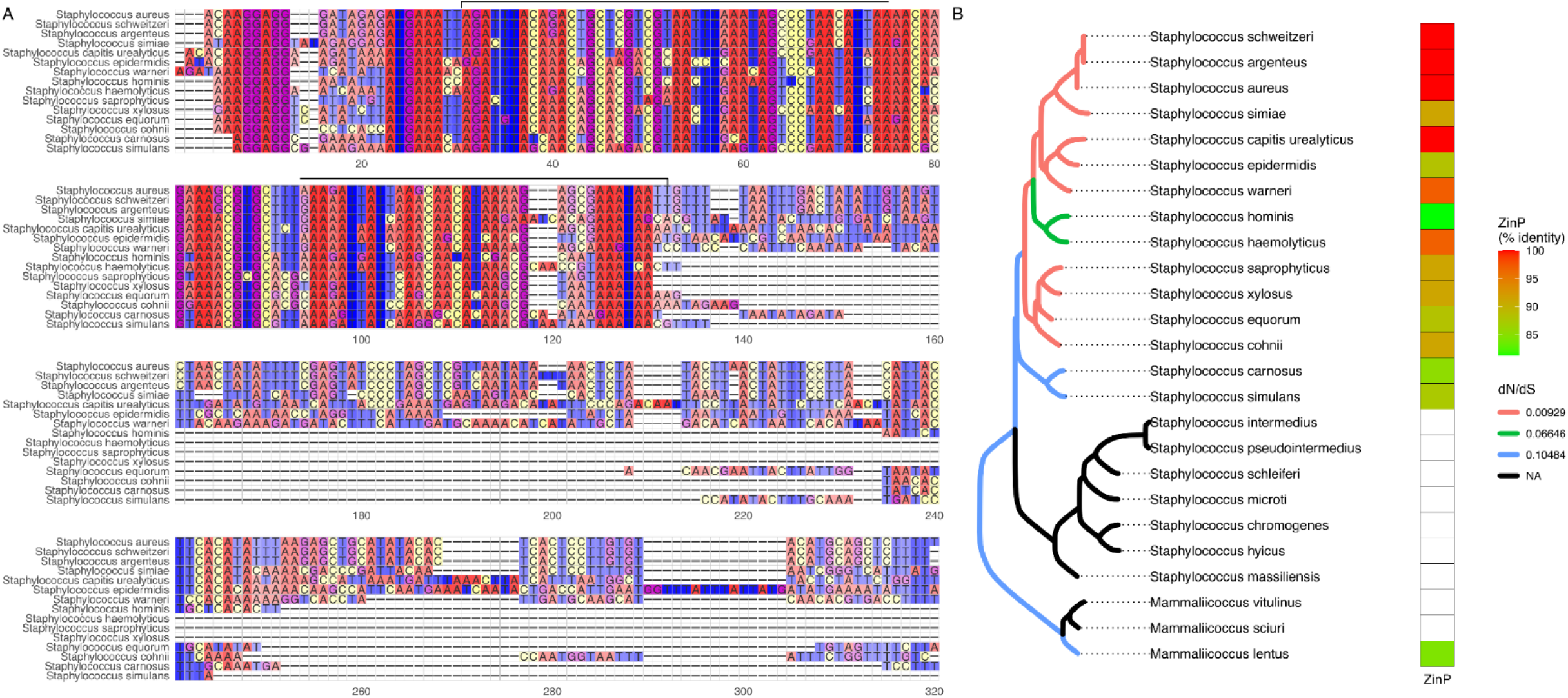
Evolution of the *zinP/S* gene. A. Sequence alignment of the region orthologous to the *zinS* locus in *S. aureus*. The upper brace indicates the ZinP coding region. B. Phylogeny of staphylococci species inferred from complete genome alignment, showing the similarity searches of the ZinP coding region. The tree branches are colored according to the inferred ratio of non-synonymous over synonymous substitutions (dN/dS), according to the model that best fit the data. See also Figure S5.

A global analysis showed that ZinP is also conserved in several organisms such as *Enterococcus* spp, *Tetragenococcus* spp, *Granulicatella* spp and *Streptococcus* spp (Figure S5), hinting that ZinP may have been acquired by horizontal gene transfer from phylogenetically distant organisms. The regulatory part of ZinS (3’) then probably emerged from the 3’UTR of ZinP. Models of codon sequence evolution revealed that ZinP is evolving under strong purifying selection, with a ratio of non-synonymous over synonymous substitutions (dN/dS) <0.11, and even <0.01 in *S. aureus*, a strong signature of protein functionality (Figure 5B).

### ZinS alters the *S. aureus* proteome in response to Zn limitation

Small RNAs typically function by base-pairings with target mRNAs, affecting their stability and/or translation, thus providing an additional layer of post-transcriptional regulation. To gain a better understanding of how ZinS shapes the staphylococcal response to Zn limitation, we performed a mass spectrometry analysis of cytoplasmic proteins in the presence and absence of Zn. We compared the cellular content between the WT strain, the Δ*zur*, Δ*zinS*, and Δ*zur* Δ*zinS* deletion mutants grown in NRPMI medium supplemented with 25 µM MnCl_2_, 1 µM FeSO_4_ ± 25 µM ZnCl_2_. The deletion of *zur* and/or *zinS* had no significant impact on growth, irrespective of the presence or absence of Zn (Figures S6B-C). The whole proteomic analysis is presented in Table S3.

We first examined the effect of Zn bioavailability by comparing the WT strain grown in the absence or presence of ZnCl_2_ (WT-vs. WT+; Figure 6A; Table S3). 69 and 38 proteins were significantly positively and negatively regulated, respectively, in the absence of Zn. As expected, the two Zn acquisition systems (AdcABC and Cnt), the non-zinc-containing paralog of the ribosomal protein S14 (RpsN2), and the putative Zn chaperone ZagA were more abundant in the absence of Zn. Concomitantly the levels of several metal-bound proteins were reduced such as the Zn-dependent alcohol dehydrogenase Adh, the copper-sensitive operon repressor CsoR, and the copper-exporting P-type ATPase A (CopA). These two examples highlight links between Zn and Cu homeostasis.

**Figure 6.**
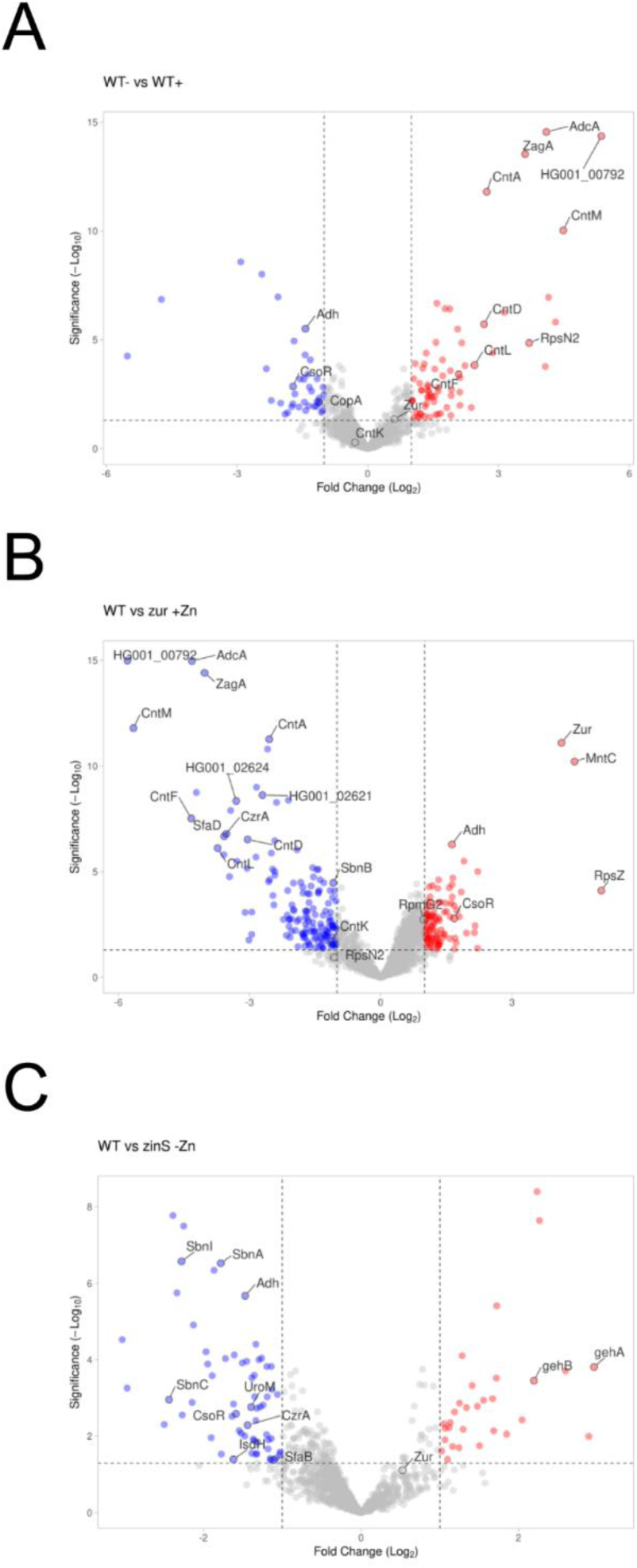
Contribution of Zur and ZinS sRNA in maintaining zinc homeostasis. Global proteomic analysis performed in WT, Δ*zur*, Δ*zinS* and Δ*zur* Δ*zinS* strains in the presence or absence of Zn. Cells were grown in NRPMI medium supplemented with 1 mM MgCl2, 100 µM CaCl2, 25 µM MnCl2, 1 µM FeSO4 ± 25 µM ZnCl2 and were harvested at OD600nm ≈ 1 (Figure S6C). A. WT strain in the absence (WT-) and in the presence (WT+) of Zn. B. WT (WT+) and Δ*zur* strain (*zur* mutant+) in the presence of Zn. C. WT (WT-) and Δ*zinS* strain (*zinS* mutant-) in the absence of Zn. Data were visualized using VolcaNoseR (Goedhart et al., 2020) and are representative of three independent experiments. The following thresholds were applied: -log10(padj)>1,3; log2FC>|1|. See Figure S6C and Table S3.

For the comparison between WT and Δ*zur* in the presence of Zn, the abundance of 250 proteins changed upon the loss of Zur: 149 were reduced and 101 were enhanced (Figure 6B; Table S3). Besides most of the previously known (e.g., ZagA, AdcA, Cnt) and *in silico* predicted targets (e.g., HG001_00792, HG001_02621-24; RegPrecise (65)) (Table S4), Zur appears to negatively regulate the zinc-responsive transcription factor CzrA and the two Fe-staphyloferrin importer systems (SbnB, SfaD). However, no clear Zur-boxes (66) were identified upstream of the related genes, suggesting an indirect regulation. In contrast, Zur seems to promote the synthesis of several metalloproteins. This includes Zn-dependent (e.g., Adh, the 30S ribosomal protein RpsZ), Cu-dependent (e.g., CsoR, the copper chaperone CsoZ), and Mn-dependent (MntC, the substrate-binding component of the Mn ABC transporter MntABC) proteins, suggesting again intricate connections between metallosystems.

We then compared the cytoplasmic protein content in the WT and Δ*zinS* backgrounds in the absence of Zn (Figure 6C; Table S3). The levels of 93 proteins were significantly impacted by the presence of ZinS (WT), among them 63 were decreased and 30 were enhanced. Interestingly, ZinS represses the synthesis of Zn-dependent (Adh, CzrA), but also Cu-dependent (CsoR), and Fe-dependent proteins (e.g., staphyloferrin A (Sfa) and B (Sbn), iron-regulated surface determinant (Isd), and uroporphyrinogen-III methyltransferase (UroM)), either directly or indirectly. Conversely, ZinS may notably induce the production of GehA and GehB lipases.

### *adh* mRNA is a main target of ZinS sRNA

To identify the targets of ZinS sRNA, we performed MS2-affinity purification coupled with RNA sequencing (MAPS) (67,68). To avoid artefactual expression using a plasmid and preserve native regulation, the MS2 sequence was chromosomally fused to the 5’ end of the *zinS* gene. Crude extracts from MS2-*zinS* and WT (control) strains were collected at an OD_600nm_ of 1 in NRPMI medium. Northern blots confirmed that the addition of the MS2 sequence did not alter ZinS levels, while control experiments confirmed that MS2-ZinS, but not ZinS, binds to an MS2 column (Figure S7A). After sequencing and data analysis, we obtained a list of 116 putative targets (FoldChange>2, P-value<0.05; Table S5). This includes mRNAs coding for Zn-dependent enzymes such as the alcohol dehydrogenase Adh, the metalloprotease Aureolysin (Aur), the L-threonine dehydrogenase (Tdh), and the molecular chaperone HslO.

To refine our analysis, we compared the MAPS results with those obtained with the CopraRNA algorithm (Table S6) (69). MAPS and CopraRNA analyses shared 24 mRNA targets (Figure S6B). We decided to validate the interaction between ZinS and seven putative targets based on their dependence on Zn (*adh*, *aur*, *tdh* and *hslO*) and/or the overlap between MAPS and CopraRNA data (*adh*, *mntR*, *isaB* and *sepF*; in the TOP20). We monitored the interaction between ZinS and putative targets using electrophoretic mobility shift assays (EMSA). We observed that ZinS sRNA forms stable complexes with *adh* (5’ part), *aur* (3’ part), *hslO* and *mntR* mRNAs *in vitro* (Figures 7A and S7). Even at high concentrations, no significant binding was observed between ZinS and *isaB*, *sepF*, and *tdh* mRNAs (Figure S7). Additionally, using the CopraRNA algorithm (Table S6), we noticed that most of the predicted pairing sites on ZinS are located from nts 190 to 230, an accessible region, being mainly single stranded and far from the *zinP* coding region (Figure 3C). This suggests that the coding and *trans*-acting activities of ZinS are dissociated.

**Figure 7.**
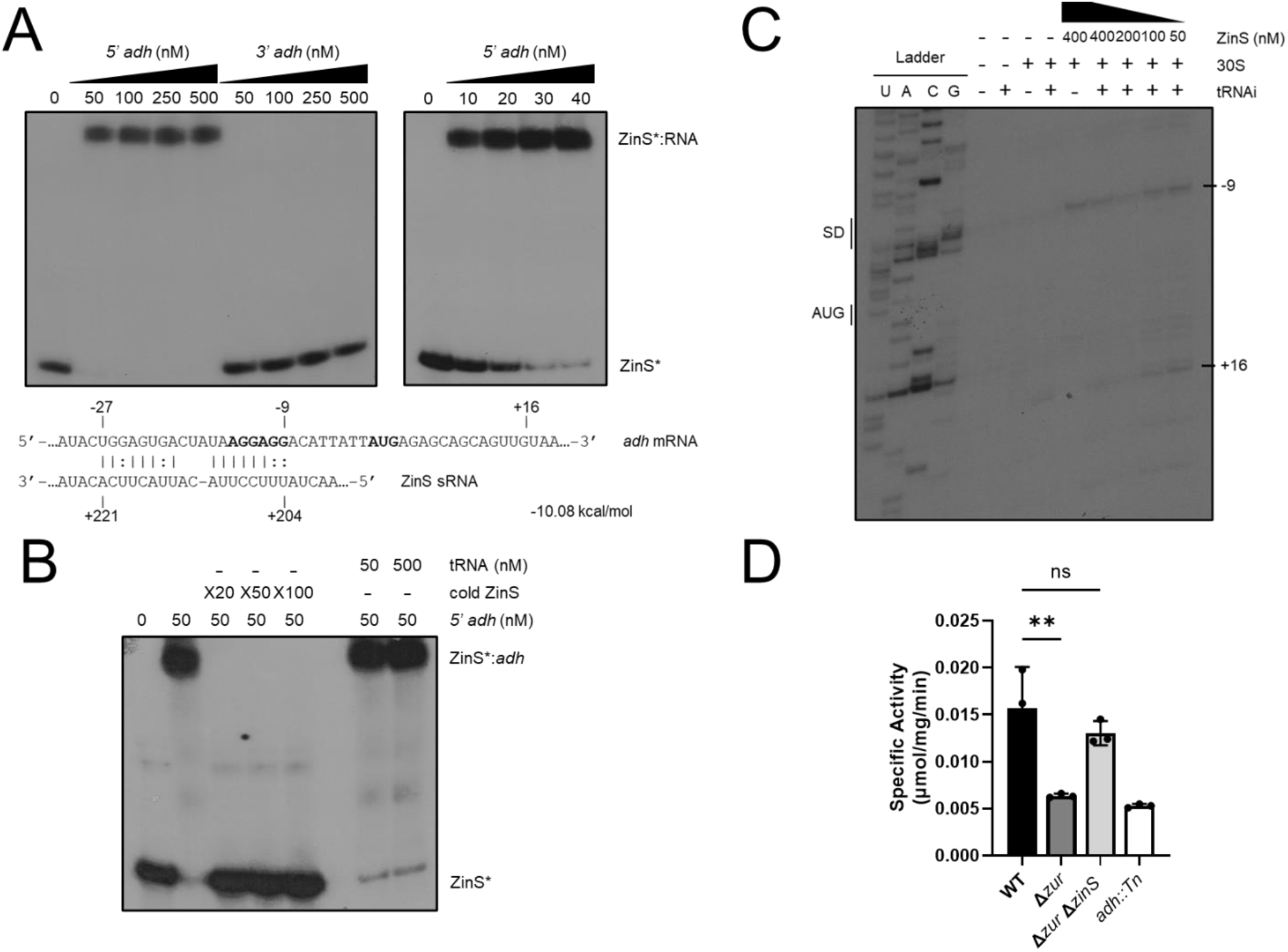
ZinS negatively regulates the Zn-dependent alcohol dehydrogenase Adh in response to zinc starvation. A. Gel retardation assays using 5’ end-radiolabeled ZinS (*) incubated with increasing concentrations of respective mRNA fragments (nM). *adh* mRNA was split into two parts, 5’*adh* (-276 to +564) and 3’*adh* (+540 to +1115). A shorter concentration range (0, 10, 20, 30, and 40 nM) was also used for 5’*adh*. Results are representative of at least two independent experiments. The ZinS:*adh* pairing site was predicted using IntaRNA software (Wright et al., 2014). The interaction energy is indicated in kcal/mol. The AUG start codon and SD sequence are in bold. (B) Gel retardation assays showing the effects of excess cold ZinS (X20, X50 and X100) or total yeast tRNA (50 and 500 nM) on the formation of the ZinS:*adh* complex (50 nM). C. Toeprinting assay monitoring the effect of ZinS on the formation of the ternary ribosomal initiation complex comprising *S. aureus* 30S subunit, initiator tRNAfMet (tRNAi) and *adh* mRNA (50 nM). Lanes 1–5: incubation controls; lanes 6–9: formation of the initiation complex in presence of increasing concentrations of ZinS (50, 100, 200, 400 nM); U, A, C, G: sequencing ladders; SD: Shine-Dalgarno sequence; AUG: start codon; -9: RT stop induced by ZinS binding; +16: toe-printing signal. D. ADH activity was quantified from cell-free lysates after culture in TSB medium for 16 hours. A one-way ANOVA with Dunnett’s multiple comparisons test was performed using Prism software (**, P-value<0.005; ns, non-significant). Data correspond to the mean of three independent experiments ± standard deviation (SD). See Figure S7.

Of the potential cytoplasmic targets, only Adh was identified by proteomics, MS2 pulldowns, and CopraRNA (Figure S7B and Table S3). Therefore, we focused on further elucidating how ZinS impacts the expression and activity of Adh. The specificity of the ZinS:*adh* interaction was further validated by competition with cold ZinS and yeast tRNA (Figure 7B). The predicted interaction site of ZinS on *adh* mRNA is located from nts -27 to -9, suggesting that ZinS sequesters the SD sequence and interferes with ribosome binding and translation initiation (Figure 7A). Therefore, we used toeprinting assays to assess the impact of ZinS binding on *adh* translation initiation (Figure 7C). We observed that ZinS inhibits the translation initiation of *adh* mRNA, as evidenced by the gradual attenuation of the +16 toe-print signal in the presence of increasing ZinS concentrations. We also noticed a second reverse transcription stop at nucleotide -9 in the presence of ZinS, which suggests that ZinS binds to the SD sequence as predicted in Figure 7A. Since ZinS prevents the recruitment of ribosomes on *adh* mRNA *in vitro*, we monitored the impact of ZinS on Adh activity *in vivo*, following spectrophotometrically the formation of NADH (Figure 7D). In the presence of ZinS, upon deletion of *zur*, we observed a significant decrease in Adh activity, similar to that observed in an *adh* mutant. The return to WT levels in a *zur zinS* double mutant further demonstrates that ZinS negatively regulates Adh synthesis and activity.

### ZinS does not alter Zn acquisition

The observation that ZinS suppresses the expression of a Zn-utilizing enzyme in response to Zn limitation is reminiscent of IsrR and RsaC, which promote survival in metal-limited environments without altering the uptake of Fe or Mn (27,28). Therefore, the impact of ZinS on cellular metal levels was evaluated using inductively coupled plasma optical emission spectroscopy (ICP-OES) following growth in NRPMI supplemented with Fe, Mn ± Zn (Figure 8). The WT and the *zinS* mutant strains accumulated comparable levels of all metals tested, including Zn, Fe and Mn. When combined with the suppression of a non-essential Zn-containing enzyme (Figure 7), these results support a role for ZinS mediating a Zn-sparing response in *S. aureus*.

**Figure 8.**
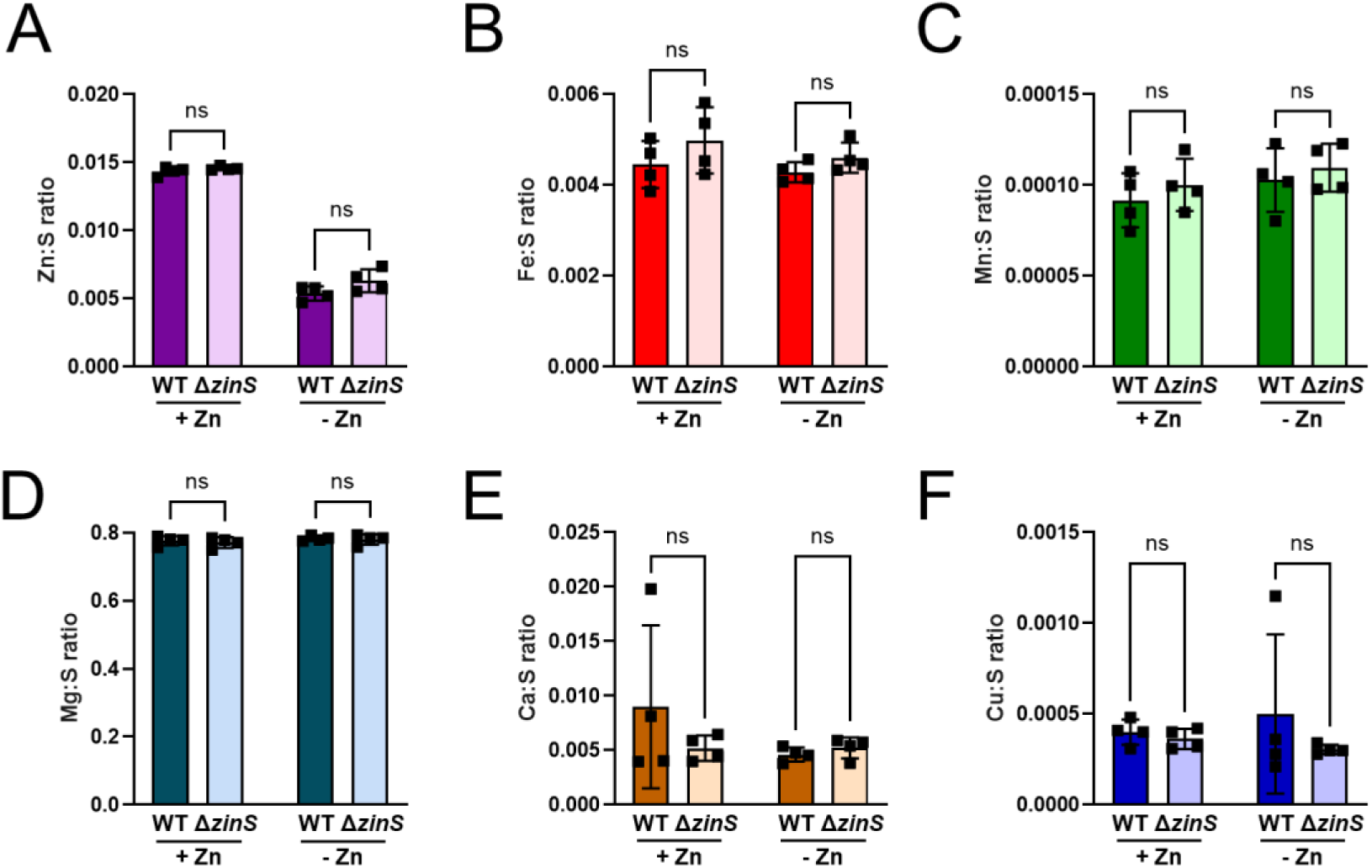
ZinS sRNA does not affect cellular accumulation of Zn. The intracellular level of (A) Zn, (B) Fe, (C) Mn, (D) Mg, (E) Ca and (F) Cu was measured in *S. aureus* wild type and Δ*zinS* using ICP-OES following growth to an OD600nm = 1 in NRPMI medium supplemented with 1 mM MgCl2, 100 µM CaCl2 ± 25 µM ZnCl2. Results were normalized to the intracellular level of sulfur (ppb). Data correspond to the mean of four independent experiments ± SD. A two-way ANOVA with Šídák’s multiple comparisons test was performed using Prism software (ns, non-significant).

## DISCUSSION

In this study, we discovered and characterized two novel sRNAs induced by zinc starvation: S1077 and ZinS (formerly RsaX20). S1077 is a 220nt-long sRNA originating from the 5’UTR of the *cnt* operon. Its synthesis is controlled by both Zur and Fur, like the staphylopine-Cnt system, whereas the 274nt-long sRNA ZinS is under the exclusive control of Zur. Besides S1077 and ZinS, 40 predicted sRNAs were activated in response to metal starvation (Figure 1; Table S2). This includes the effector of the *agr* quorum-sensing system RNAIII (70), confirming recent findings showing its enhanced expression in the presence of calprotectin (71). Conversely, 42 sRNAs were repressed such as RsaI and SprC. The catabolite control protein A (CcpA)-repressed sRNA RsaI is a key actor of the metabolic switch occurring during glycolysis. It primarily regulates glucose uptake, catabolism, and biofilm formation, potentially providing a link between metal homeostasis and carbon metabolism (72). The importance of sRNAs in fine tuning metabolism in response to metal availability is evidenced by ZinS suppression of Adh, which contributes to fermentative metabolism. Interestingly, SprC sRNA reduces the synthesis of the Zn efflux pump CzrB (73). SprC was previously described as involved in resistance to oxidative stress, immune escape, and attenuation of *S. aureus* virulence (74). However, its role in Zn homeostasis remains unclear, as the SprC-mediated regulation of CzrB is detrimental to *S. aureus* in the presence of zinc toxicity (73), and is only known to be negatively controlled by SarA, which does not respond to zinc (75).

The role of sRNAs was previously reported for Fe homeostasis in a number of other microbes, where the tandem pair of the Fur protein and RyhB-like sRNAs critically modulate the response to Fe limitation (3). In *S. aureus*, Fe, Mn, and Zn homeostasis are all controlled by a tandem of transcription factor and sRNA: Fur-IsrR (26,27), MntR-RsaC (25) and Zur-ZinS. A common characteristic of the Fe and Mn regulatory circuits is that transcription factors control transport systems, while sRNAs contribute to metal-sparing by suppressing the synthesis of metal-utilizing proteins. For example, RyhB-like sRNAs (e.g., RyhB in *E. coli*, PrrF in *Pseudomonas aeruginosa*, FsrA in *Bacillus subtilis* and IsrR in *S. aureus*) inhibit the synthesis of many Fe-containing enzymes, notably involved in the TCA cycle, such as aconitase and succinate dehydrogenase (27,76–80). RyhB-like and RsaC sRNAs also share a similar strategy to spare their cognate metal: both interfere with the synthesis of metal-utilizing superoxide dismutases (25,77,78). Upon Zn starvation, ZinS negatively regulates Zn-containing proteins, such as the alcohol dehydrogenase Adh. This suggests that this sRNA also activates a metal-sparing response. Additionally, ZinS could have a broader role in metal homeostasis and virulence via the regulation of the Mn-dependent transcription factor MntR and the metalloprotease Aureolysin, involved in immune evasion (81). Although ZinS is the first Zur-regulated sRNA described in bacteria, it seems likely that other species utilize sRNAs to adapt to Zn limitation and activate a Zn-sparing response, given the conservation of this regulatory logic for other metals and the universal essentiality of Zn.

In this study, we also expanded the regulon of Zur. In addition to previously known targets (Table S4), we identified Zur-boxes upstream of the (i) HG001_02621-02624 operon, coding for a putative Zn-binding GTPase (COG0523 subfamily), a putative iron transporter and a putative oxidoreductase, and (ii) HG001_00792, coding for a putative pyridine nucleotide-disulfide oxidoreductase. Their synthesis is induced by CP treatment (82,83), confirming our proteomic analyses (Figure 6B). Remarkably, we found that any perturbation of Zn homeostasis (Figure 6) also affects key proteins involved in other metal-related adaptive responses, such as metal importers (Mn: MntABC; Fe: Sbn, Sfa, Isd), metal exporters (Cu: CopA), and transcription factors (Cu: CsoR; Zn: CzrA). Intricate connections between metallosystems have already been described in bacteria. As mentioned above, the metallophore staphylopine is produced via Zur and Fur dual regulation in *S. aureus* (12,16). However, the Cnt system also drives copper uptake, triggering the copper stress response (84). In *Streptococcus agalactiae* a link between Cu and Zn homeostasis has also been observed (85). Additionally, several studies highlighted close interactions between Fe and Mn homeostasis (86–88). Overall, these interconnected regulatory networks allow bacterial cells to optimally respond to encountered stresses by integrating multiple environmental signals.

ZinS is a dual-function sRNA, which codes for a peptide renamed ZinP. This is not an isolated example in bacteria, as other regulatory RNAs share this feature (e.g., RNAIII encoded *hld* in *S. aureus* and RybA encodes *mntS* in *E. coli*) (89,90). Although, the function of ZinP remains unknown, its expression pattern and localization near metal transporters (Figure S4) suggests that ZinP also plays a role in metal homeostasis. When adapting to metal-restricted environments, bacteria induce the expression of metal uptake systems, but they also reduce the expression of efflux pumps to prevent secretion of a now limited resource. While transcriptional and translational regulation prevent the production of new efflux pumps, they do not inactivate previously produced proteins. During the transition from high to low Mn, the Mn efflux pump MntP and the sRNA-encoded peptide MntS are simultaneously present in *E. coli* cells (64). Although MntP synthesis is tightly repressed at the transcriptional level, existing MntP pumps are still active at the membrane. Therefore, as part of a Mn-sparing response, MntS inhibits Mn efflux by interacting directly with MntP. As the ZinP structure is reminiscent of the MntS structure (Figure 4A), it is tempting to speculate that ZinP has a similar role in *S. aureus*, potentially by inhibiting Zn efflux via CzrB.

Gene duplication and gain-of-function mutations are key mechanisms of bacterial adaptive evolution. For example, the amplification of genes encoding iron transporters confers a selective advantage to overcome nutritional immunity (91). In *S. aureus*, the superoxide dismutase SodM was obtained by duplication of *sodA* and then neofunctionalized to utilize either Mn or Fe, a key advantage for resisting manganese starvation (92,93). Remarkably, we showed that the regulatory part of ZinS likely emerged from the 3’UTR of ZinP after horizontal gene transfer from phylogenetically distant organisms. This is reminiscent of the 3’UTR-derived sRNA RsaC, released from the *mntABC* polycistronic transcript, encoding the main Mn importer in *S. aureus* (25). The fact that both were exclusively acquired by *S. aureus* and closely related strains indicates that these organisms are under strong selective pressure to acquire additional mechanisms to survive metal limitation. Remarkably, the acquisition of RsaC allows the integration of the two superoxide dismutases, SodA and SodM, into a new regulatory circuit (25). To preserve functional detoxification of reactive oxygen species (ROS) during Mn starvation, RsaC specifically inhibits the synthesis of the Mn-dependent SodA and indirectly promotes the switch to the cambialistic SodM. Hence, both ZinS and RsaC are striking illustrations that 3’UTRs serve as an important reservoir of unrestricted RNA sequence for the evolution of regulatory elements. In line with our study, recent observations have shown that large 3’ UTRs evolved rapidly to confer species specificity and, in turn, contribute to bacterial diversity (94). We previously hypothesized that the presence of 5′→3′ exoribonucleases RNases J1/J2 might present a significant obstacle to the evolution of 3′UTR sRNAs in Gram-positive bacteria (95). However, this barrier can be bypassed by using various strategies to protect the 5’ end from RNase attack. Our studies showed that the 3’UTR of ZinP is not released as an independent 3’UTR sRNA, and consequently retains the protective 5’PPP. Although RsaC sRNA is released from its precursor by RNase III cleavage, resulting in the presence of a 5’P, the presence of a long 5’ secondary structure certainly reduces its accessibility to RNases (25). While numerous 3’UTR-encoded sRNAs have been described in Gram-negative bacteria (96,97), it now becomes clear that this evolutionary strategy is also widely used by Gram-positive bacteria (25,98–100).

In conclusion, this study showed that staphylococci obtain a messenger RNA via horizontal transfer and, in a second step, neofunctionalized its 3’UTR to gain regulatory functions. This perfectly illustrates how bacteria acquire new regulatory elements and integrate them into new regulatory circuits in response to strong selective pressure, such as metal restriction imposed by the host immune system.

## DATA AVAILABILITY

Both raw and processed data obtained from RNA sequencing are available in the Gene Expression Omnibus database under the accession numbers GSE296223 (sRNA-seq) and GSE296224 (MAPS). The mass spectrometric data were deposited to the ProteomeXchange Consortium via the PRIDE partner repository with the dataset identifier PXD065909.

## SUPPLEMENTARY DATA STATEMENT

Supplementary Data are available at *NAR* Online.

## ACKNOWLEDGMENTS

The authors would like to thank Dr. Alejandro Toledo-Arana and all team members for helpful discussions.

## AUTHOR CONTRIBUTIONS STATEMENT

Conceptualization, T.E.K. and D.L.; Formal analysis, J.N.R., J.J.T., K.J.W., J.Y.D., D.L.; Methodology, M.C., G.R., V.M., J.C., J.J.T., K.J.W., J.Y.D., T.E.K, and D.L.; Validation, J.N.R, V.M., J.C., J.J.T., K.J.W., J.M.B., J.Y.D., T.E.K, and D.L.; Investigation, M.C., S.P., H.R., J.N.R., G.R., H.M.L, M.P.K., R.M., M.B., V.M., J.C., J.J.T., J.Y.D., and D.L.; Writing – Original Draft, T.E.K. and D.L.; Writing – Review & Editing, J.N.R., P.R., J.J.T., K.J.W., J.M.B., J.Y.D, T.E.K., and D.L.; Visualization, M.C., J.N.R., G.R., M.P.K., J.C., J.J.T., J.Y.D., and D.L.; Supervision, S.M., K.J.W., J.M.B., T.E.K., and D.L.; Project Administration, D.L.; Resources, V.M., J.C., K.J.W., J.J.T., J.Y.D., T.E.K, and D.L.; Funding Acquisition, P.R., J.J.T., K.J.W., J.M.B., T.E.K, and D.L.

## FUNDING

D.L. and P.R. were supported by the Agence Nationale de la Recherche (ANR-20-CE12-0021 and ANR-23-CE12-0041-01). D.L. and T.E.K were supported by Thomas Jefferson Fund, a program of FACE Foundation launched in collaboration with the French Embassy. D.L., J.C., S.M., and P.R. were supported by the Interdisciplinary Thematic Institute IMCBio+, as part of the ITI 2021-2028 program of the University of Strasbourg, CNRS and Inserm, IdEx Unistra (ANR-10-IDEX-0002), SFRI-STRAT’US (ANR-20-SFRI-0012), and EUR IMCBio (ANR-17-EURE-0023) under the framework of the French Investments of the France 2030 Program. Work in the laboratory of T.E.K. was supported by grants from the NIH (R01AI179695 and AI155611) and by the University of Iowa’s Year 2 P3 Strategic Initiatives Program through funding received for the project entitled “High Impact Hiring Initiative (HIHI): A Program to Strategically Recruit and Retain Talented Faculty”. The Boyd lab was supported by National Science Foundation award 1750624 and USDA MRF project NE−2248. K.J.W. was supported by a MAESTRO grant from the National Science Center (NCN), Poland (2021/42/A/NZ1/00214). J.J.T. was supported by grants from the National Health and Medical Research Council (GNT2028572) and Australian Research Council (DP220101938).

## CONFLICT OF INTEREST DISCLOSURE

The authors declare no competing interests.

## Notes

### Competing Interest Statement

The authors have declared no competing interest.

