## Supplementary data for "A Zur-dependent regulatory RNA involved in maintaining zinc homeostasis in *Staphylococcus aureus*"

### SUPPLEMENTARY MATERIALS AND METHODS

#### Conservation analysis

Nucleotide BLAST (NCBI) was used to compare the sequences of the sRNAs (S1077 and ZinS). EFI tools were used to study the conservation of the ZinP peptide (1). First, we used the Enzyme Similarity Tool (EST), which allows the comparison and visualization of the sequence similarity between enzymes. EST generates sequence similarity networks (SSNs), which are used as input for the Genome Neighborhood Tool (GNT). Basically, GNT retrieves information about the genes in the vicinity of the input enzymes in different genomes and highlights conserved gene clusters.

#### MS2-affinity purification coupled with RNA sequencing (MAPS)

The MS2 sequence was fused to the 5' end of the *zinS* gene using the pMAD-*MS2-zinS* vector and homologous recombination (Table S1B). Consequently, the *MS2-zinS* construct is regulated by the endogenous promoter. The MS2 addition was predicted to have no significant impact on the ZinS secondary structure using mfold algorithm. To induce zinc starvation, we used NRPMI medium supplemented with 1 mM $MgCl_2$ and 100 µM $CaCl_2$. Crude extracts from MS2-*zinS* and WT (control) strains were collected at an $OD_{600nm}$ = 1. MS2-affinity purifications were conducted in duplicate, following the procedure described by Mercier et al. (2021). Bacterial cells were resuspended in Lysis Buffer (20 mM Tris-HCl pH 8, 150 mM KCl, 1 mM MgCl2, 1 mM DTT) and broken using the FastPrep apparatus (MP Biomedicals). Crude extracts were loaded on a MS2-MBP coated column and purified by affinity chromatography. Eluted RNA were extracted with phenol:chloroform:isoamyl alcohol and subsequent ethanol precipitation. DNase I (0.1 U/ml, Roche) treatment was performed 30 min at 37°C. RNA quantity and quality were assessed using the 2100 Bioanalyzer (Agilent). Ribosomal RNA was depleted from RNA samples using the QIAseq FastSelect rRNA removal kit (QIAGEN), and cDNA libraries were prepared with the NEBNext Ultra II directional RNA library prep kit for Illumina (NEB). Sequencing of cDNA libraries was performed on the NextSeq 2000 system (Illumina). RNA-seq analysis was carried out according to the method outlined by Lalaouna et al. (2018) (2).

#### *In silico* prediction of ZinS targets

A list of putative targets has been obtained using CopraRNA (default parameters) and the sequence of *zinS* genes from *S. aureus* subsp. *aureus* HG001 isolate RN1 (NZ_CP018205), *S. aureus* subsp. *aureus* USA300_FPR3757 (NC_007793), *S. aureus* subsp. *aureus* str. Newman (NC_009641), *S. aureus* subsp. *aureus* N315 (NC_002745), *Staphylococcus argenteus* 58113 (NZ_AP018562) and *Staphylococcus schweitzeri* NCTC13712 (NZ_LR134304).

**Proteomic analysis**

WT, Δ*zur*, Δ*zinS* and Δ*zur* Δ*zinS* cells grown in NRPMI medium supplemented with 1 mM $MgCl_2$, 100 µM $CaCl_2$, 25 µM $MnCl_2$, 1 µM $FeSO_4$, ± 25 µM $ZnCl_2$ were harvested at $OD_{600nm} \approx 1$. Cell pellets were washed 5 times with cold DPBS, resuspended in 200 µL lysis buffer (7 M urea, 2 M thiourea, 4% CHAPS, and 20 mM Tris-HCl pH 7.6, 67 U of Pierce Universal Nuclease (ThermoFisher Scientific) and 1 mM $CaCl_2$), and lysed using Fastprep homogenizer. The Bradford assay was used to determine protein concentrations. For mass spectrometry analysis, 10 µg of purified proteins were precipitated with 0.1 M ammonium acetate in 100% methanol (5 volumes, -20°C). Proteins were reduced (5 mM Dithiothreitol, 95°C, 10 min), alkylated (10 mM Iodoacetamide, room temperature, 20 min) and digested overnight with 1:50 (v/v) of sequencing-grade trypsin (Promega). Peptide mixtures were analyzed by nanoLC-MS/MS on a reversed phase nanoElute 2 coupled to a TIMS-TOF Pro 2 mass spectrometer (Bruker Daltonik Gmbh) using a data-dependent acquisition (DDA). Peptides were separated on the integrated emitter column IonOpticks Aurora Elite (25 cm × 75 µM, 1.7 µm particle size and 120 Å pore size; AUR3–15075C18-CSI) with a 70 min gradient. Proteins were identified via comparison to a homemade *S. aureus* protein database (2623 sequences), analyzed using Mascot algorithm (version 2.8, Matrix Science) with a decoy strategy. Mass error was set to 20 ppm for precursor ions and 30 ppm for fragment ions. Oxidation (M), carbamidomethylation (C), acethylation (protein Nter) and phosphorylation (S, T and Y) were considered as variable modifications. Data were then imported into Proline v2.0 software (3), proteins were validated on Mascot pretty rank equal to 1, a Mascot score threshold set at 25 and 1% FDR on both peptide spectrum matches (PSM score) and protein sets (Protein Set score) and MS1 eXtracted Ion Chromatograms (XIC) were used to quantify each protein. An exhaustive map alignment followed by a median ratio normalisation of the intensities was computed on Proline. A 50s cross assignment was carried out within groups only. Prostar was used for the statistical analyses of the intensities. Partially observed values (POV) and values missing in the entire condition (MEC) were imputed with det quantile 1%. A LIMMA statistical test with a log (FC) threshold of 1 and a Benjamini-Hochberg correction were used to generate log2(FC) and adjusted p-values. Data were visualized using VolcaNoseR (4). Results are representative of three independent experiments.

#### Growth monitoring assays

Freshly streaked bacterial colonies were used to inoculate overnight cultures incubated at 37°C with constant shaking (180 rpm). Cultures were diluted ($OD_{600nm}$ = 0.05) in fresh media before being placed in a 96-well flat-bottomed plate and then incubated at 37°C with constant agitation for 20 hours. Growth was measured at $OD_{600nm}$ in a microplate reader Multiskan SkyHigh multiplate spectrophotometer (ThermoFisher). At least three independent experiments were performed. Data were analyzed and visualized using Prism software.

#### Determination of the 5' end status of S1077

To determine if the 5' end of S1077 is mono- or tri-phosphorylated, we used Terminator 5'-Phosphate-Dependent Exonuclease (Biosearch Technologies) according to manufacturer's protocol. Total RNA (10 $\mu$g) was incubated with Terminator Exonuclease (1U) and Terminator 1X Reaction Buffer A at 30°C for 1 h. RNA were then purified by phenol extraction and ethanol precipitation.

#### dRNA-seq and Term-seq

dRNA-seq and Term-seq data were obtained from Mediati et al. (2022) and processed as previously. The data are available under GEO accession GSE158830.

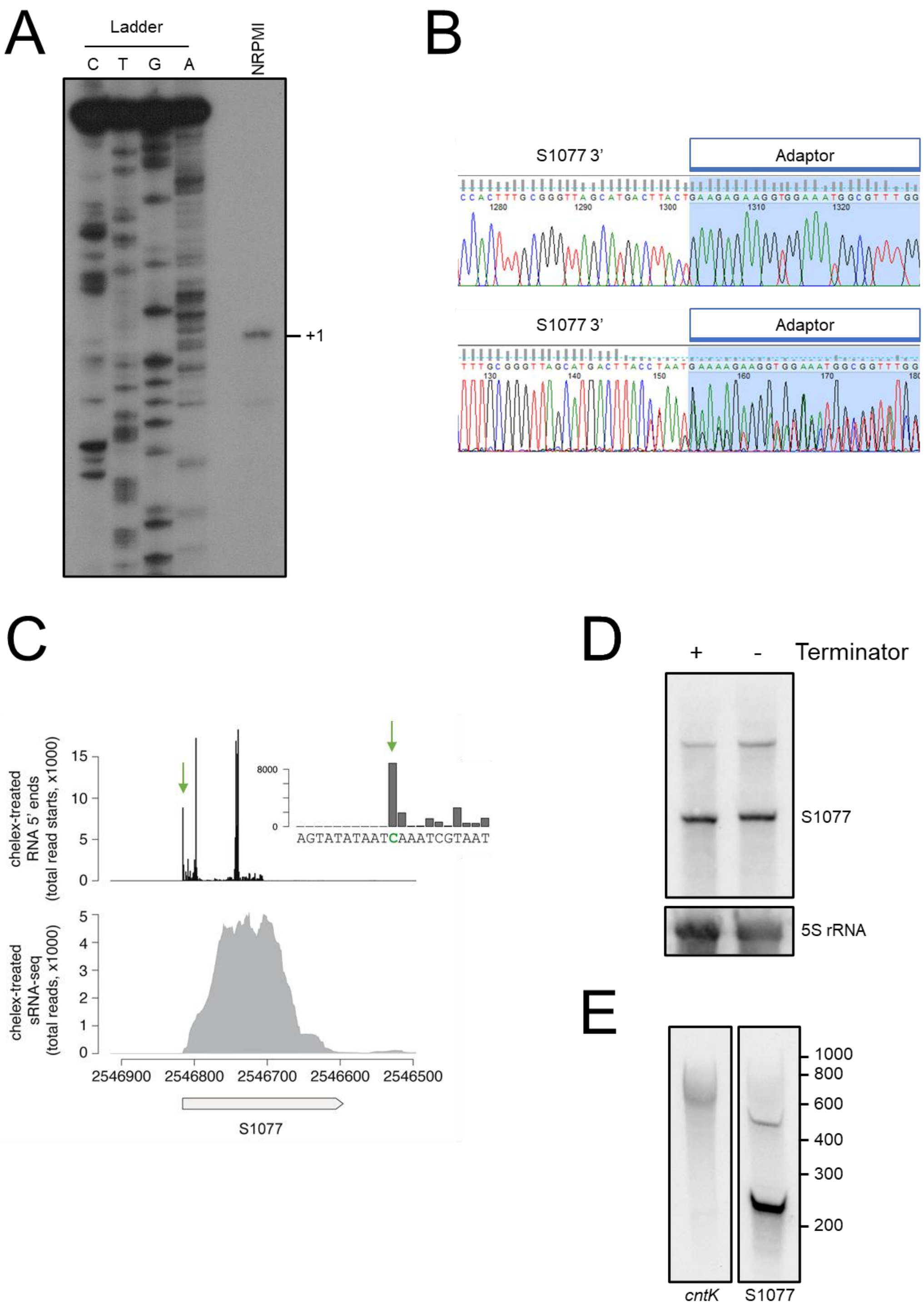


**Figure S1. Determination of S1077 boundaries.** A. Primer extension assay performed on total RNA extracted in NRPMI medium supplemented with 1 mM $MgCl_2$ and 100 µM $CaCl_2$ ($OD_{600nm}$ = 1) using another 5'-radiolabelled oligonucleotide that is complementary to the S1077 sequence. This second primer is located at nucleotides +166 to +186 (from the +1 of S1077). The determine transcription start of S1077 is indicated by +1. G, A, C, T: sequencing ladders. Data are representative of two independent experiments. B. Two representative chromatograms showing the sequence obtained after sequencing of cDNA fragments obtained by 3'RACE and visualized with FinchTV. In blue is the adaptor sequence. Note that there is a reproducible C>T mutation for the first identified 3'end. C. End mapping of S1077 in *S. aureus* strain HG001. The boundaries of S1077 were mapped in strain HG001 using sRNA-seq of chelex-treated cultures. The inset shows the sequence at the 10 nucleotides around the 5' end peak and genomic coordinate. The grey box indicates position of S1077 in the HG001 genome. D. Northern blot analysis of total RNA extracted from cells grown in NRPMI medium supplemented with 1 mM $MgCl_2$, 100 µM $CaCl_2$ and 25 µM $MnCl_2$, treated with the Terminator 5'-Phosphate-Dependent Exonuclease (Biosearch technologies)**.** 5S rRNA was used as a control, being protected from exonuclease attack. Results are representative of two independent experiments. E. Northern blot analysis using S1077 and *cntK* specific DIG probes. Total RNA was extracted from HG001 WT cells grown in NRPMI medium supplemented with 1 mM $MgCl_2$, 100 µM $CaCl_2$. Cells were harvested at $OD_{600nm}$ = 1. RNA migrated on an acrylamide gel along with a RiboRuler Low Range RNA Ladder (Thermo scientific). Related to Figure 2.

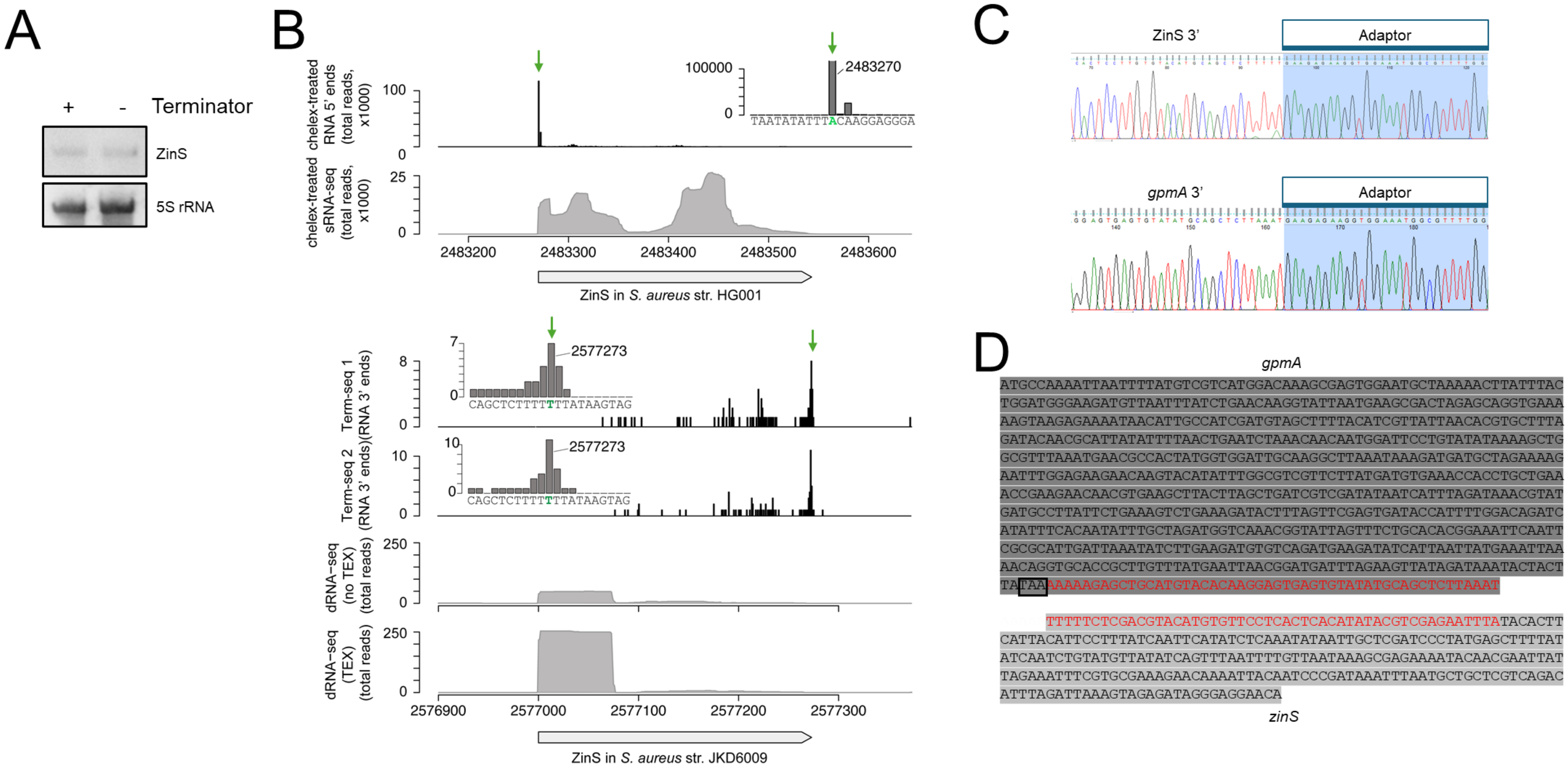


**Figure S2. The 3'end of ZinS sRNA and *gpmA* mRNA are overlapping.** A. Northern blot analysis of total RNA extracted from Δ*zur* cells grown in RPMI + 1% CA medium, treated with the Terminator 5'-Phosphate-Dependent Exonuclease (Biosearch technologies). 5S rRNA was used as a negative control, being protected from exonuclease attack. Results are representative of two independent experiments. B. End mapping of ZinS in *S. aureus* strains HG001 and JKD6009. (Top panels) The boundaries of ZinS were mapped in strain HG001 using sRNA-seq of chelex-treated cultures. The inset shows the sequence at the 10 nucleotides around the 5' end peak and genomic coordinate. The grey box indicates position of ZinS in the HG001 genome. (Bottom panels) The boundaries of ZinS were mapped in strain JKD6009 using dRNA-seq to identify RNA 5' ends (with or without terminator exonuclease, TEX). RNA 3' ends were mapped using Term-seq (replicates indicated by 1 and 2). The inset shows the sequence at the 10 nucleotides around the 3' end peak and genomic coordinate. Data originate from Mediati et al. (2022). C. Representative chromatogram showing the sequence obtained after sequencing of cDNA fragments obtained by 3'RACE (*zinS* and *gpmA*) and visualized with FinchTV. In blue is the adaptor sequence. See also Figure 2B. D. The overlapping sequence is shown in red. *gpmA* (CDS and 3'UTR) and *zinS* (opposite strand) sequences are shown in dark and light gray, respectively. The stop codon of *gpmA* is boxed. Related to Figure 3.

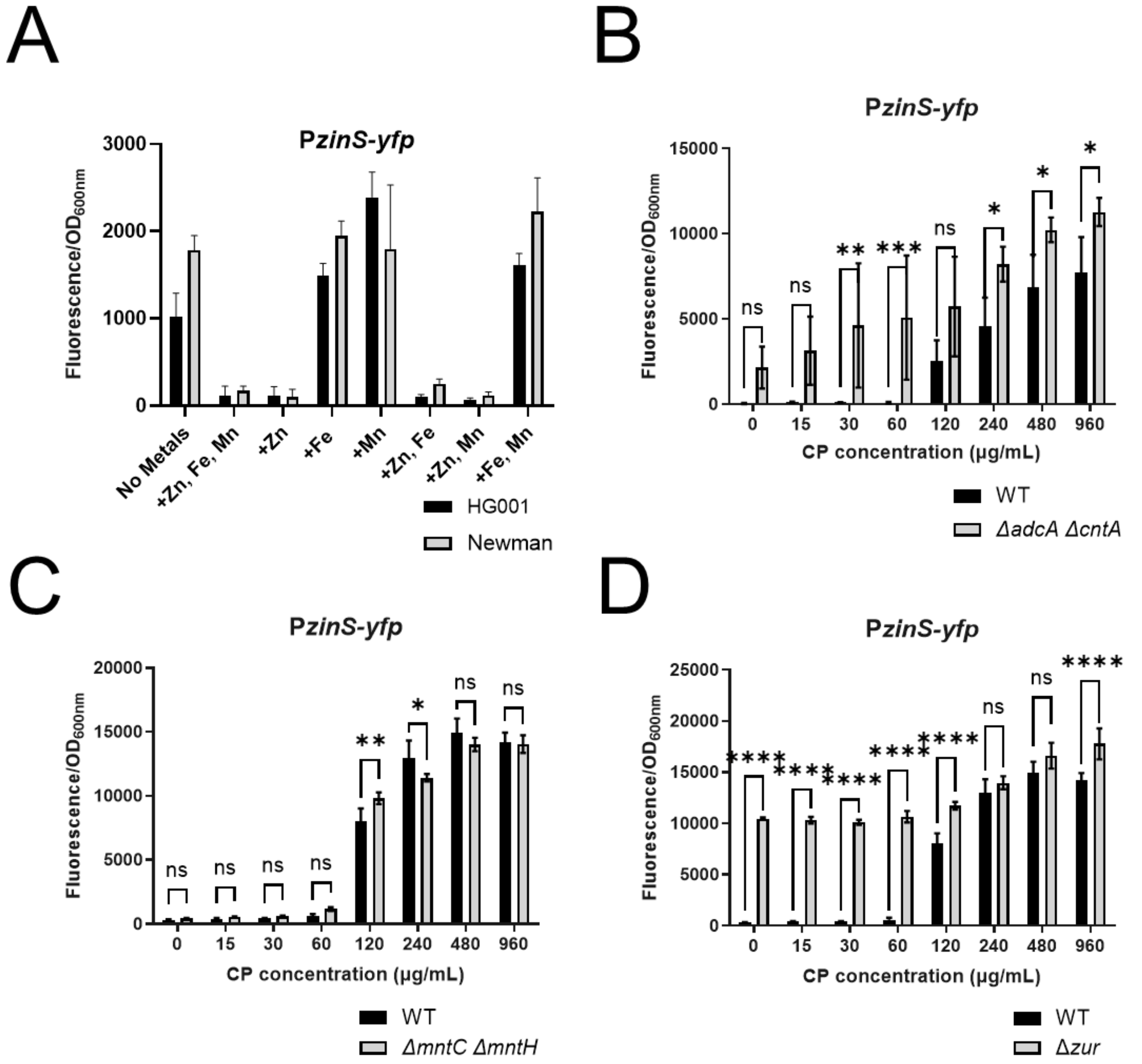


**Figure S3. The transcriptional activity of *zinS* increases in response to Zn limitation.** A. S. *aureus* HG001 (black) and Newman (gray) strains, containing P*zinS-yfp* transcriptional fusion, were grown in NRPMI medium supplemented with indicated metals. The ratio fluorescence/OD$_{600nm}$ was determined after 8h of growth. Data correspond to the mean of three independent experiments ± SD. B-D. *S. aureus* Newman strains, containing P*zinS-yfp* transcriptional fusion, were grown in TSB medium supplemented with 1 µM $MnCl_2$ and 1 µM $ZnCl_2$ in the presence of increasing concentrations of calprotectin (CP; 0 to 960 µg/mL): (B) WT (black) and Δ*adcA* Δ*cntA* (gray), (C) WT (black) and Δ*mntC* Δ*mntH* (gray), and (D) WT (black) and Δ*zur* (gray). The ratio fluorescence/OD$_{600nm}$ was determined after 8h of growth. Data correspond to the mean of three independent experiments ± SD. A two-way ANOVA with Šídák's multiple comparisons test was performed using Prism software (*, P-value<0.05, **, P-value<0.005, ***, P-value<0.0005, ****, P-value<0.0001; ns, non-significant). Related to Figure 3.

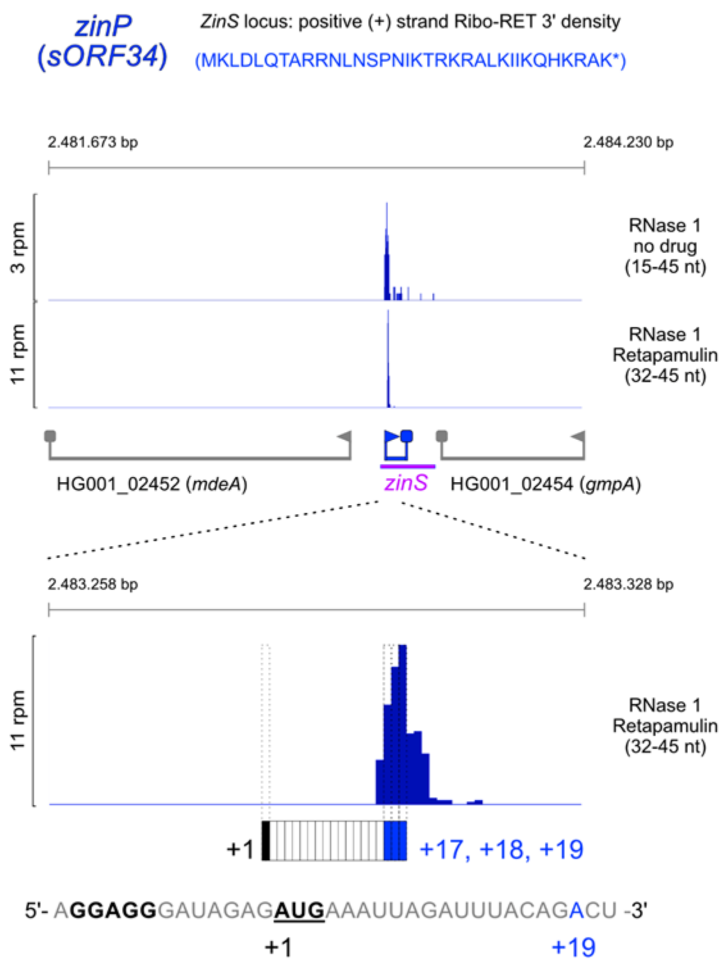


**Figure S4. Ribosome profiling indicates efficient translation of ZinP *in vivo*.** RNAse 1 Ribo-seq and Ribo-Ret reads from published sequencing data (Kohl et al., 2025), highligthing ZinP ribosomal footprints (RPFs). The enriched density at the ZinP start site in the Ribo-Ret data (in response to Retapamulin treatment) strongly indicates efficient translation initiation. Respective positive strand 3' end density profiles of the *zinS* locus are shown and the treatment condition (no drug vs. Retapamulin) as well as the selected RPF size for display are indicated accordingly. Note that translation initiation generates larger RPF size in the dataset and is particularly enriched, selecting RPFs of 32 to 45 nt length. A zoomed in view corresponding to ribosomal density of the ZinP translation initiation peak is presented together with the sequence designating its ribosomal binding site, SD-motif and start codon. The read 3' end density clusters around the expected distance from the start codon (+17 nt) as analyzed by published metagene analysis for this dataset (Kohl et al., 2025). All screenshots were captured on a custom R-studio riboseq density visualization app. Related to Figure 4.

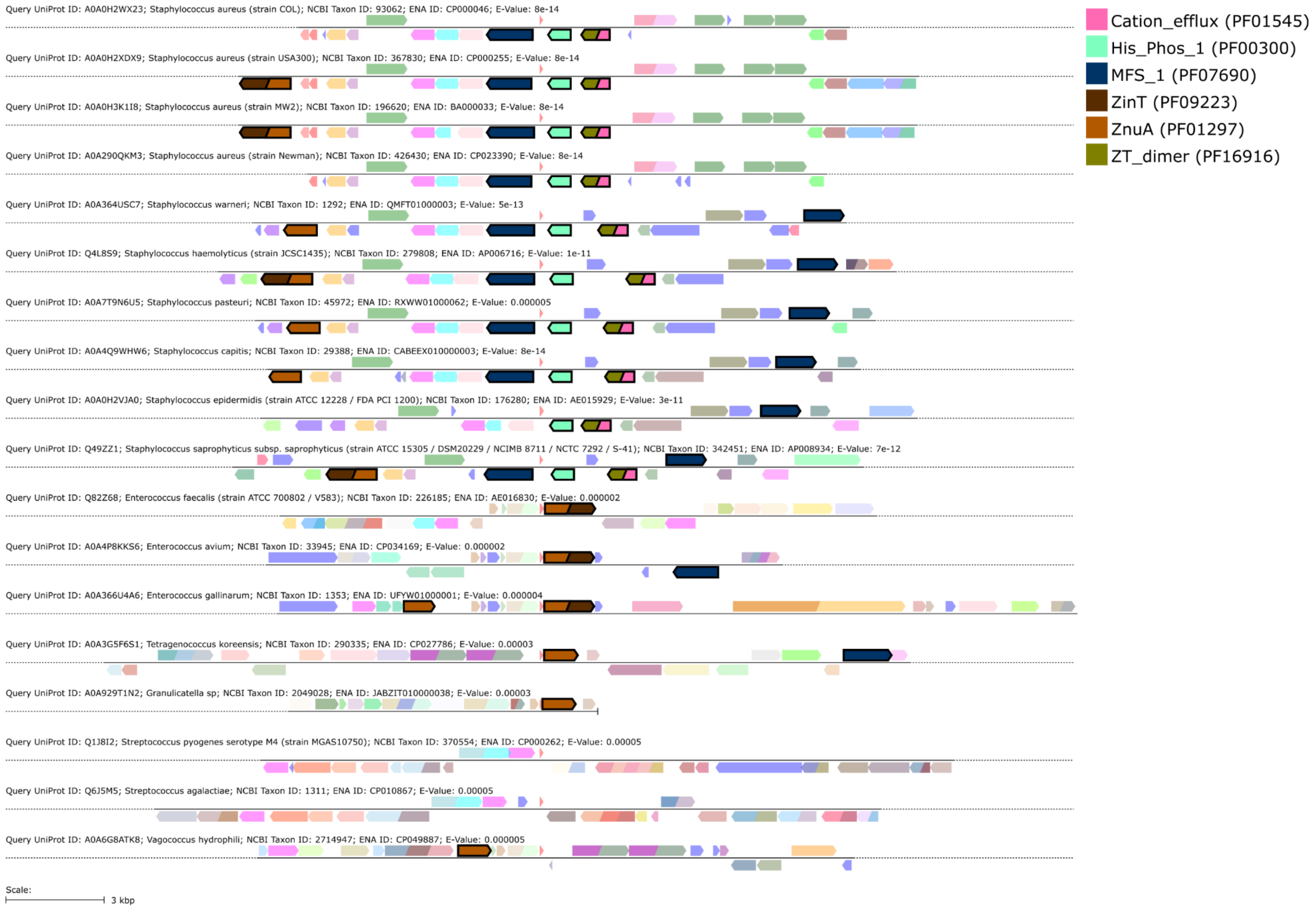


**Figure S5. Synteny analysis of the small peptide encoded by ZinS.** The EFI - Enzyme Similarity Tool (Oberg et al., 2023) was used to perform a BLAST search using *zinP* sequence. This analysis generates a sequence similarity network (SSN) used by EFI-Genome Neighborhood Tool to place protein families and superfamilies into a genomic context. Relevant protein domains with their Pfam entry are indicated in the upper right-hand corner. ZinP is indicated by a red triangle in the middle. Related to Figure 5.

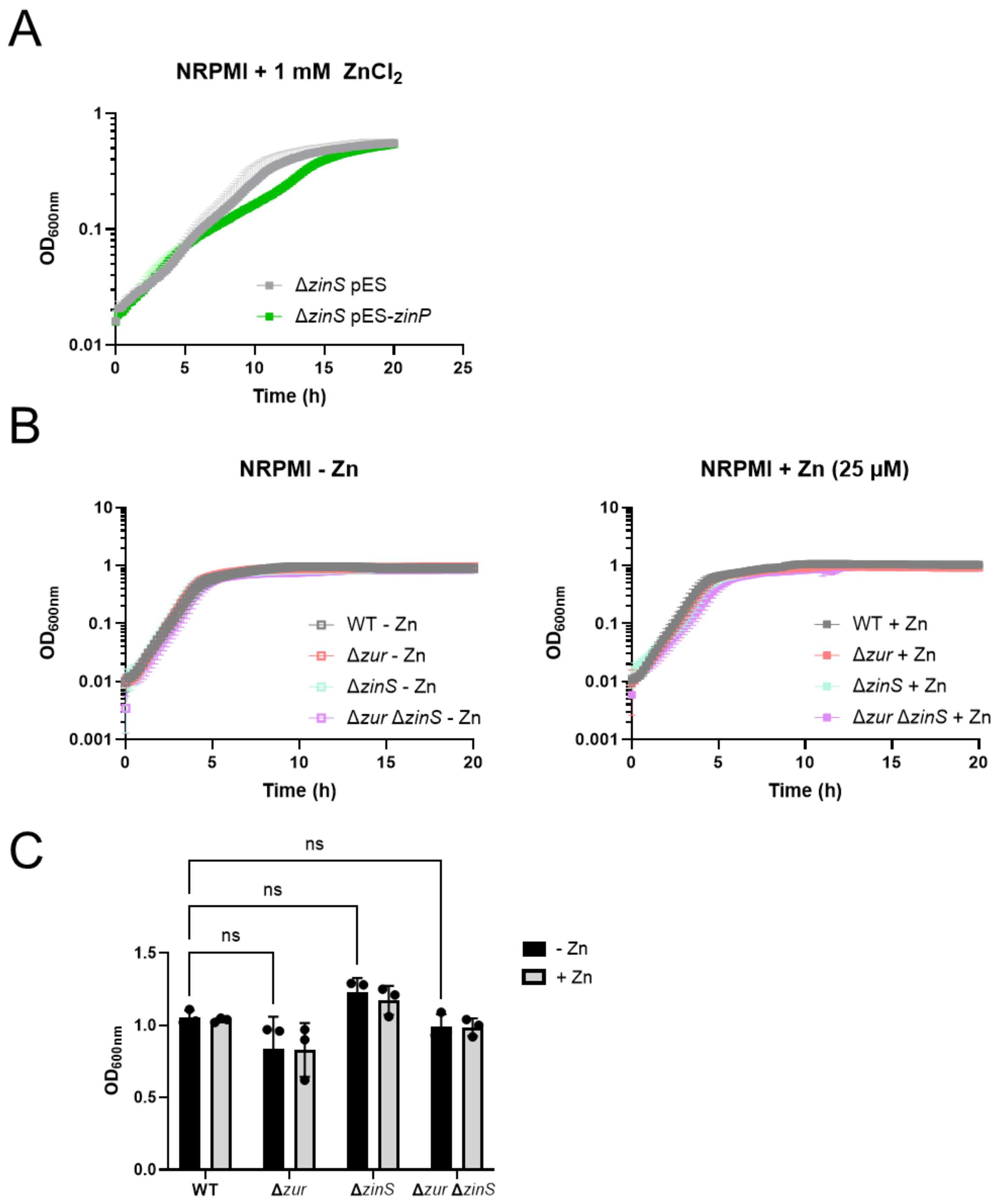


**Figure S6. Impact of Zur, ZinS, and ZinP on *S. aureus* growth.** A. Growth monitoring (20h) of the Δ*zinS* strain containing the empty vector or a pES derivative plasmid which enables the strong and constitutive expression of *zinP*. Cells were grown in NRPMI medium supplemented with 1 mM $MgCl_2$,100 µM $CaCl_2$, 25 µM $MnCl_2$, 1 µM $FeSO_4$ and 1 mM $ZnCl_2$. Data correspond to the mean of three independent experiments ± SD. B. Growth monitoring (20h) of WT, Δ*zur*, Δ*zinS*, and Δ*zur* Δ*zinS* strains grown in NRPMI medium supplemented with 1 mM $MgCl_2$,100 µM $CaCl_2$, 25 µM $MnCl_2$, 1 µM $FeSO_4$ ± 25 µM $ZnCl_2$. Data correspond to the mean of four independent experiments ± SD. C. The $OD_{600nm}$ was measured when the cells were collected for proteomic analysis. A two-way ANOVA with Šídák's multiple comparisons test was performed using Prism software (ns, non-significant). Related to Figure 6.

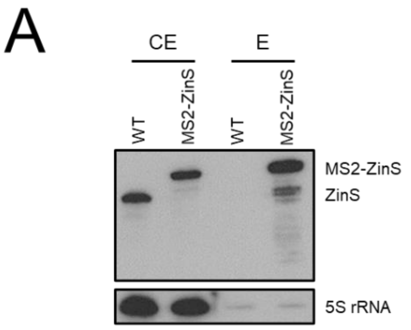


B

| Names | Total | mRNAs |
|---|---|---|
| CopraRNA, MAPS and MS | 1 | *adh* |
| CopraRNA and MAPS | 23 | HG001_01038 HG001_01262 HG001_00224 HG001_02319 *lcpC* *mntR* aroGH isaB HG001_02547 HG001_02156 HG001_02356 *sepF* HG001_00635 *queH* HG001_00549 *sle1* *ccpA* *galM* HG001_02317 *mvaA* *folK* HG001_02376 HG001_02478 |

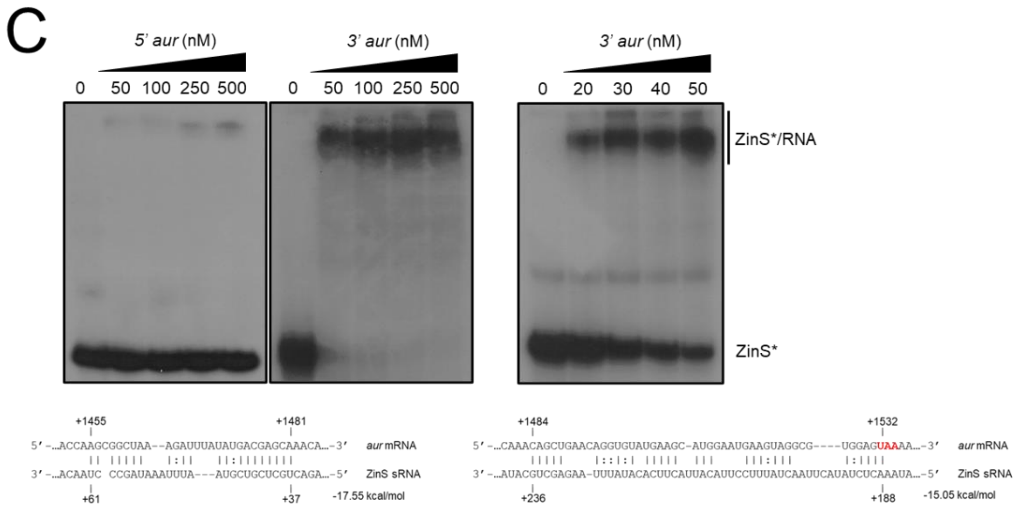


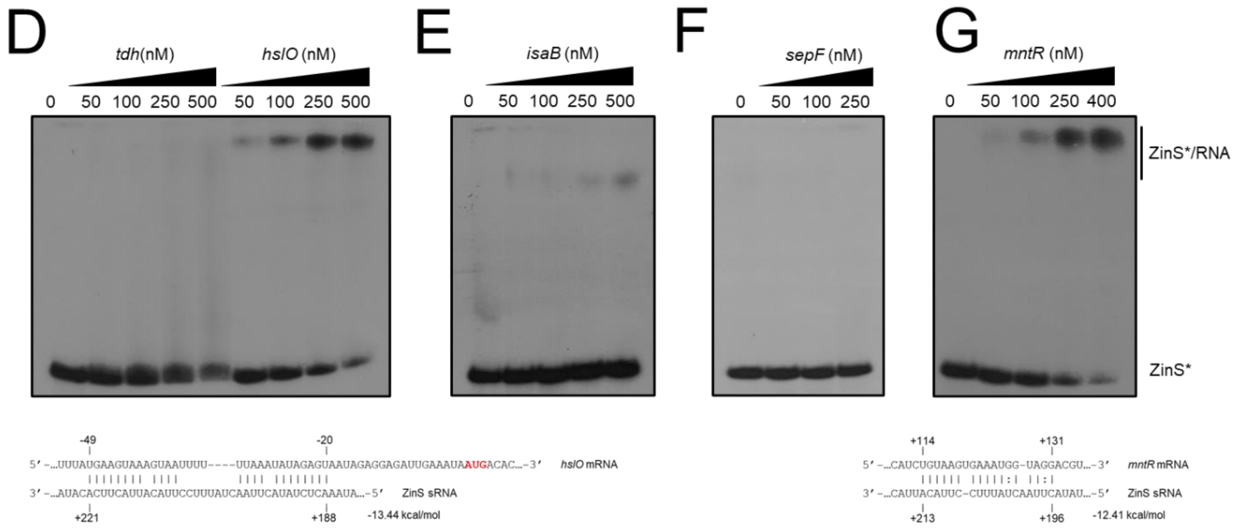


**Figure S7. Identification of the ZinS targetome through a combination of experimental and computational approaches.** A. Northern blot analysis of ZinS and MS2-ZinS RNA levels after MS2-affinity purification samples using a ZinS-specific DIG probe. WT (Control) and MS2-*zinS* cells were grown in NRPMI medium supplemented with 1 mM $MgCl_2$ and 100 µM $CaCl_2$ to induce metal starvation. The impact of MS2 was evaluated using mfold algorithm. Cells were collected at $OD_{600nm}$ = 1. 5S rRNA was used as loading control. Crude extract (CE); Elution (E). B. Comparison of candidate mRNAs/proteins identified by MAPS (Table S5), CopraRNA (Table S6) or mass spectrometry (MS; Table S3). The putative mRNA targets that have been tested by EMSA are shown in red. (C-G) Gel retardation assays using ZinS sRNA and putative mRNA targets. 5' end-radiolabeled ZinS (*) was incubated with increasing concentrations of selected mRNAs. C. *aur* mRNA was split into two parts, 5'*aur* (-40 to +745) and 3'*aur* (+718 to 1660). A shorter concentration range (0, 20, 30, 40, and 50 nM) was also used for 3'*aur*. D. *tdh* mRNA (-67 to +662). E. *isaB* (full-length; -39 to +589). F. *sepF* (-200 to +568). G. *mntR* mRNA (full-length; -20 to +713). The ZinS:mRNA pairing sites were predicted using IntaRNA when complex formation was observed (Wright et al., 2014). The two most likely predictions are indicated for 3'*aur*. The interaction energy is indicated in kcal/mol. The AUG start codon or STOP codon are in red. Results are representative of at least two independent experiments. Related to Table S5.
